# OsbZIP62/OsFD7, a functional ortholog of Flowering Locus D (FD), regulates floral transition and panicle development in rice

**DOI:** 10.1101/2021.03.09.434543

**Authors:** Amarjot Kaur, Aashima Nijhawan, Mahesh Yadav, Jitendra P. Khurana

**Author notes:** Correspondence: Prof. Jitendra P. Khurana. While giving finishing touches to this article, Dr. Aashima Nijhawan left for heavenly abode at a very young age. We dedicate this article to the memory of Dr. Aashima Nijhawan and will always fondly remember her as a person and also with the zeal and passion she worked all through her stay in the laboratory.

## Abstract

We have characterized a rice bZIP protein coding gene *OsbZIP62*/*OsFD7* that expresses preferentially in SAM and during early panicle developmental stages in comparison to other OsFDs characterised till date. Surprisingly, unlike OsFD1, OsFD7 interacts directly and more efficiently with OsFTLs; the interaction is strongest with OsFTL1 followed by Hd3a and RFT1, as confirmed by FLIM-FRET analysis. Also, OsFD7 is phosphorylated at its C-terminal end by OsCDPK41 and OsCDPK49 *in vitro* and this phosphorylated moiety is recognized by OsGF14 proteins. *OsFD7* RNAi transgenics were late flowering; the transcript levels of some floral meristem identity genes (e.g. *OsMADS14*, *OsMADS15* and *OsMADS18*) were also down-regulated. It was quite interesting to note that these RNAi lines exhibited dense panicle morphology with increase in the number of primary and secondary branches resulting in longer panicles and more seeds probably due to downregulation of *Sepallata* (*SEP*) family genes. In comparison to other FD-like proteins characterized thus far from rice, it appears that OsFD7 may have undergone diversification during evolution resulting in the acquisition of newer functions and thus playing dual role in floral transition and panicle development in rice.

**Highlight:** OsbZIP62/OsFD7 interacts with major flowering regulators participating in the processes of floral transition as well as panicle and floral organ development.

## INTRODUCTION

The transition from vegetative to reproductive development in plants is a complex process initiating the conversion of shoot apical meristem (SAM) into floral meristem. Analyses of a large number of *Arabidopsis* mutants and natural variants have contributed immensely in understanding this complex regulatory network (Coupland, 1995; Koornneef and Peeters, 1997; Adrian *et al*., 2009; Huijser and Schmid, 2011). Whereas *Arabidopsis* represents a dicot system, among the monocots, rice has been the system of choice for studying the complex floral induction and evocation pathways. In *Arabidopsis*, under inductive photoperiods, Flowering locus T protein (FT; mobile flowering signal called as florigen), homologous to the mammalian phosphatidylethanolamine binding protein (PEBP) family (Chardon and Damerval, 2005; Corbesier *et al*., 2007), expressing in leaves moves *via* phloem to SAM where it interacts with FD, a basic-leucine zipper (bZIP) transcription factor. In fact, FD protein is first phosphorylated at its C-terminal end (L-X-R/K-X-X-S/T) by certain calcium dependent protein kinases (CDPKs), which in turn is recognized by intracellular receptor G-box factor 14-3-3-like protein (GF14), a phospho-protein binding adaptor protein. The three-way protein interaction between phosphorylated FD, GF14 and FT results in the establishment of a functional complex at SAM activating floral meristem identity genes like *Leafy* (*LFY*), *Suppressor of Overexpression of Constans 1* (*SOC1*), *Fruitfull* (*FUL*) and *Cauliflower* (*CAL*), *Apetala1* (*AP1*), orchestrating floral organ initiation and development (Abe *et al*., 2005; Wigge *et al*., 2005; Kawamoto *et al*., 2015). Essentially a similar pathway operates in rice wherein OsFD1 interacts with Hd3a/OsFTL-2 via OsGF14, forming a tripartite complex called FAC that initiates floral transition by modulating the expression of OsMADS15 and other MADS-box transcription factors (Taoka *et al*., 2011). In rice, 13 *FT-like* genes have been identified. Among these, *Hd3a/OsFTL-2* is the closest ortholog of *AtFT* (Kojima *et al*., 2002; Chardon and Damerwal, 2005), followed by *OsFTL-1* and *OsFTL-3/RFT1*, and they promote flowering in rice (Yano *et al*., 2001; Izawa *et al*., 2002).

A large number of bZIP protein coding genes were first identified in *Arabidopsis* by Jakoby *et al*. (2002) and subsequently in rice by our group (Nijhawan *et al*. 2008) and some of them have been functionally validated to perform diverse functions in plants (CorrÍa *et al*., 2008; Nijhawan *et al*., 2008; Burman *et al*., 2018; Dr^ge-Laser *et al*., 2018). Based on sequence analyses of these bZIP proteins and homology search, Tsuji *et al*. (2013) identified at least five more OsFDs, namely, OsFD2, OsFD3, OsFD4, OsFD5, and OsFD6. Whereas OsFD1 and OsFD4 specifically regulate flowering (Taoka *et al*., 2011; Cerise *et al*., 2020), OsFD2 plays important role in leaf development (Tsuji *et al*., 2013). Similarly, the bZIP proteins Liguleless2 (LG2) and Delayed Flowering1 (DLF1) in maize play important role in reproductive transition (Walsh and Freeling, 1999; Muszynski *et al*., 2006). Another FD-like protein in potato interacts with StSP6A and GF14-3-3 proteins, forming a triprotein complex, just like FAC, and triggers tuber formation (Teo *et al*., 2017).

The availability of finished quality sequence of the rice genome (IRGSP, 2005) opened up avenues for functional genomic studies worldwide (Vij *et al*., 2006; The Rice Annotation Project, 2008). We also embarked on the microarray-based whole genome expression analyses in rice to identify genes expressing preferentially during floral transition and panicle development in rice (e.g. Jain *et al*., 2007, 2008, Arora *et al*., 2007; Agarwal *et al*., 2007; Kapoor *et al*., 2008; Nijhawan *et al*., 2008; Sharma *et al*., 2012). In this study, we have made an attempt to elucidate the role of *OsbZIP62*/*OsFD7* protein coding gene in both floral transition and panicle development, since, unlike other *OsFD* genes characterized till date, it preferentially expresses in SAM and at very early stages of panicle development (Nijhawan *et al*., 2008). We have named this OsbZIP as OsFD7. Moreover, OsFD7 interacts directly with OsFTL1 and OsGF14s providing evidence for its role in floral transition. We have also shown interaction of OsFD7 with OsCDPKs, phosphorylating its C-terminal end. We also provided evidence for its possible involvement in panicle and floral organ development. This study has thus revealed that, in comparison to the roles ascribed to other rice FD or FD-like proteins in rice, OsFD7 has acquired novel functions in reproductive development in rice during the course of evolution.

## MATERIALS AND METHODS

### Gene amplification and plasmid construction for plant transformation

*OsFTL1* and *OsbZIP62*/*OsFD7* cds (Fig. S1A) were amplified from KOME clones obtained from NIAS (National Institute of Agrobiological Sciences) (*OsFTL1*-AK072979; *OsFD7*-AK110915). *Hd3a, RFT1*, *OsRCN1*, *OsGF14b,c,d* and *OsCDPK41,49* genes were amplified from different cDNA samples. Deletion constructs of *OsFD7* were prepared by overlap extension PCR. All the mutant constructs of *OsFD7* were generated by incorporating mutation in the reverse primer sequences. For generating *OsFD7* RNAi transgenics, a 312 bp region (953 bp to 1260 bp fragment of *OsFD7* cDNA, amplified using KOME clone with accession number AK110915, with CACC at 5’ end) was cloned in pENTR-D-TOPO vector using pENTR^TM^ Directional TOPO® cloning kit (Invitrogen Inc. USA). This fragment was mobilized to pANDA vector (Miki and Shimamoto, 2004) using LR clonase Enzyme mix II kit (Invitrogen Inc., USA) following manufacturer’s protocol. Rice transgenics in PB1 (*Oryza sativa* ssp *indica* Pusa Basmati 1) background were generated according to Mohanty *et al*. (1999). Transgenic nature of these rice transgenics was confirmed by isolating genomic DNA and performing PCR amplification of *HPTII* gene.

### Expression profiling by real-time PCR analysis

Rice tissue samples were harvested from plants grown in the fields at Indian Agricultural Research Institute, New Delhi, and their RNA isolated using TRIzol reagent following manufacturer’s protocol (Chomczynski and Sacchi, 1987); these samples represented root, mature leaf, Y-leaf, flag leaf, SAM: up to 0.5 mm, P1-1: 0.5-2 mm, P1-2: 2-5 mm, P1-3: 5-10 mm, P1: 0-3 cm, P2: 3-5 cm, P3: 5-10 cm, P4: 10-15 cm, P5: 15-22 cm, P6: 22-30 cm (*see* Jain *et al*., 2007; Nijhawan *et al*., 2008). cDNA was synthesized from 1 µg RNA using High capacity cDNA archive kit (Applied Biosystems, USA). Real-time PCR was performed using LightCycler© 480 real-time system as per manufacturer’s protocol (Roche, Germany).

### Phenotypic characterization of rice RNAi transgenics and expression analysis of floral meristem and organ identity genes

All the transgenics and control plants were grown under natural conditions in a containment net house during the normal rice-growing period (from June to September), every year. Flowering time was determined based on days required for the emergence of panicle (i.e. heading date). Many morphological characteristics were recorded, including panicle length, number of primary and secondary branches, seed harvest (total number of seeds per panicle), % sterility (number of sterile florets per panicle), seed dimensions: length and width (Image J software) and 100 grain weight. These data have been represented graphically and statistical significance calculated using student’s *t*-test. In addition, transcript accumulation of floral meristem and organ identity genes was quantitated by performing real-time PCR of SAM, P1-1, P1-2 and P1-3 stage tissues of RNAi transgenics and vector control (VC) plants.

### Total protein extraction and western analysis

Total protein was extracted from different tissues using protein extraction buffer [200 mM Tris pH 8.0, 100 mM NaCl, 10 mM Na_2_EDTA, 1% protease inhibitor cocktail (Roche), 10 mM DTT, 0.05% Tween20 and 5% glycerol]. The tissue was ground in a pre-chilled mortar and pestle using liquid nitrogen. To this homogenized sample, 500 µL of buffer was added and it was allowed to thaw on ice. The protein sample was transferred to 1.5 mL MCT and centrifuged for 20 min at 13,000 rpm, 4°C. The supernatant was collected in fresh MCT and quantified using Bradford reagent (Bradford, 1976). These protein samples were stored at -80°C. For determining protein level of OsFD7 by western analysis, 5 µg of protein extracts were utilized. Polyclonal OsFD7 antibody generated in rabbit was used at a dilution of 1:20,000 + 1% w/v BSA and secondary anti-rabbit antibody at a dilution of 1:50,000 + 1% w/v BSA for detecting protein levels by western blotting. Actin monoclonal antibody (Thermo Fisher Scientific) was utilized for detecting actin protein as a loading control in all the samples tested.

### Yeast transactivation and interaction studies

For yeast assay, different gene constructs were cloned in pGADT7 and pGBKT7 vectors (Clontech Laboratories Inc., USA). For three-way interaction analysis, CDS of *OsFD7* was cloned in MCSI and CDS of *OsGF14b,c* was cloned in MCSII of pBRIDGE vector. Primer sequences are given in Table S1. Small scale yeast transformations were carried out in Y2H Gold (for pGADT7 and pGBKT7 constructs) and AH109 (for pBRIDGE constructs) yeast strains following manufacturer’s protocol. The transformation mixes were plated on SD/-W (pGBKT7 construct), SD/-LW (for pGADT7 and pGBKT7 constructs) and SD/-LMW (for pGADT7 and pBRIDGE constructs). Primary and secondary cultures were initiated at 30°C and drop assay carried out by placing 10 µL drops of serial dilutions (10^-1^, 10^-2^, 10^-3^ and 10^-4^) prepared in autoclaved MQ water, on different selection media supplemented with 1-5 mM 3-AT along with respective control media (SD/-HW and SD/-W for transactivation assay using pGBKT7 vector constructs; SD/-HLW and SD/-LW for yeast one-to-one interaction assay using pGBKT7 and pGADT7 vector constructs; SD/-HLMW and SD/-LMW for yeast three-hybrid interaction assay using pBRIDGE and pGADT7 vector constructs). The plates were kept at 30°C for 3-5 days.

### Particle bombardment of onion peel cells

The CDS sequences of *OsFD7*, *OsFTL1*, *Hd3a*, *OsGF14b* and *OsCDPK41,49* were amplified and cloned in: pSITE-3CA, pSITE-1NB vectors for localization and FLIM-FRET analysis; pSITE-3CA-nYFPC1 as well as pSITE-3CA-cYFPC1 for BiFC analysis via gateway technology as described earlier. Particle bombardment of onion peel cells was performed using biolistics particle delivery system PDS-1000/He (Bio-Rad, USA) as per the protocol of Lee *et al*. (2008) with the following set parameters: 1300 psi of He pressure and 6 cm of distance from launch assembly. The plates were kept in dark at 28°C. After 16 h incubation, YFP fluorescence was visualized under the Leica® TCS, SP5 confocal microscope. Appropriate negative controls for BiFC were chosen according to the guidelines given by Kudla and Bock (2016). FLIM-FRET analysis was performed according to De Los Santos *et al*. (2015) and Long *et al*. (2018) on Leica SP8-FALCON FLIM microscope. All these experiments were repeated thrice for every combination with at least five cells analyzed in each experiment.

### Protein induction and purification

Different gene constructs used in this study were cloned in pET28a(+), pET29a(+) (Novagen) and pGEX4T1 (GE) vectors (refer Supporting Table S1 for primer sequences). These clones were transformed into BL21-(DE3)-RIL or Rosetta-gami^TM^ competent cells. Induced proteins were sonicated and their supernatant fractions utilized for further experiments. Protein concentration of different induced samples was determined by performing Bradford assay (Bradford, 1976). Supernatant fraction of OsFD7 was purified using Ni-NTA column (Qiagen) and used for antibody production (Radiant Research, Bangalore, India). Antibody was raised against the whole protein.

### *In vitro* pull-down assay

Supernatant fractions of the induced proteins were employed for all the GST pull-down experiments. For pull-down assay, 100 µL of GST-Sepharose beads were washed thoroughly with MQ water followed by the addition of 500 µL of equilibration buffer (100 mM Tris-Cl pH 8.0, protease inhibitor cocktail tablet, 10 μg). The proteins were allowed to adsorb for 2 h at 4°C with constant shaking. After μ adsorption, beads were washed with 1 mL of equilibration buffer at least ten times to remove unbound protein from the GST-Sepharose slurry. After washing, 200 μ L of equilibration buffer containing 0.5% NP40 and μ incubated at 4°C for 16 h with constant shaking. Beads were washed at least five times with 500 μL of equilibration buffer containing 0.5% NP40 and was finally suspended in 100 μ same buffer. Western blotting was performed using anti His-tag (Cat no. H1029-5ML, Sigma, USA) as well as with anti GST antibody (Cat no. G7781-2ML, Sigma, USA). GST protein was used as a control for all the pull-down experiments. For detecting protein interactions of OsFD7 with OsCDPKs in the presence or absence of calcium, 1 mM CaCl_2_ and 1 mM of EGTA was added, respectively, to the equilibration buffer. Similar procedure was followed for pull down assay between S-tagged and His-tagged proteins. His-tagged protein was adsorbed onto Ni-NTA slurry and S-tagged protein sample was incubated with this adsorbed Ni-NTA slurry. Western blotting was performed using anti S-tag (Cat no. 71549-3, Novagen) and anti His-tag antibody to detect protein-protein interaction.

### Electrophoretic mobility shift assay

G/C-box and a sequence lacking G/G-box (Non-GB/CB oligo) were used for performing gel shift assay as described by Taoka *et al*. (2011). The gels were stained with LabSafe^TM^ Nucleic acid stain (Cat. #786-409) and then destained in MQ water. See Table S1 for oligo sequences.

### *In vivo* phosphorylation assay

A 30 µg aliquot of total protein isolated from SAM, P1-1, P1-2 and P1-3 stage tissues each was incubated in a buffer (25 mM Tris-HCl pH 7.5, 10 mM EGTA, 10 mM EDTA, 150 mM NaCl, 10% glycerol, 0.1% Triton X-100, 1 mM DTT, 1 mM Na_3_VO_4_, 1 mM NaF and protease inhibitor cocktail) with 2 µl of OsFD7 primary antibody. This mixture was rotated constantly at 4° C for 2 h and then centrifuged for 10 min at 15,000 g, 4°C. The pellet obtained was washed thrice with the buffer and finally resuspended in 20 µl of buffer. 5X loading dye was added and the samples boiled for 5-10 min. These protein samples were resolved on SDS-PAGE and western analysis performed using anti-phosphoserine/phosphothreonine antibody (Matsushita *et al*., 2013; Cheng *et al*., 2014).

### Protein dephosphorylation assay

A 5 µg aliquot of total protein isolated from SAM tissue was utilized for immunoprecipitating OsFD7 protein with 1 µl of OsFD7 primary antibody in a MCT using the protocol mentioned in the above section. The immunoprecipitated protein was incubated with λ protein phosphatase using manufacturer’s protocol (Cat. # P0753S, NEB). The treated samples were resolved on SDS-PAGE and western blotting performed using OsFD7 antibody.

### *In vitro* kinase assay

250 ng purified bacterially expressed OsFD7 protein was incubated with 500 ng purified fraction of OsCDPK49 in a kinase buffer (40 mM Tris-Cl pH 7.5, 15 mM MgCl_2_, 0.5 mM CaCl_2_, 2 mM DTT, 500 µM ATP) with or without calcium (EGTA 0.5 mM) at 30°C for 30 min (Lambda protein phosphatase - 0.5 µl + 1X MnCl_2_ -30°C for 30 min). The reaction was terminated by adding 5X loading dye and the samples boiled for 5-10 min. The protein samples were resolved by SDS-PAGE and western analysis performed using anti-phosphoserine/phosphothreonine antibody.

### Phylogenetic and motif analysis

Protein sequences of different FD and FD-like genes were retrieved from Phytozome v10.1 web server (http://www.phytozome.net/). All these protein sequences were aligned using the MAFFT server (Multiple Alignment with Fast Fourier Transform; http://mafft.cbrc.jp/alignment/software/). MAFFT aligned sequence file was uploaded on PhyML 3.0 web server (http://www.phylogeny.fr/) and phylogenetic tree was generated. The file generated was visualized using FigTree (http://tree.bio.ed.ac.uk/software/figtree/). SMART domain analysis software (Schultz *et al*., 1998) was used to predict the domains present in OsFD7. Additional conserved motifs in OsFD7 were determined using SALAD database with parameters such as “maximum number of motifs to find” set at 10 and “expect threshold” value set to 1e-2 (Mihara *et al*., 2010).

### Structural determination, interaction and localization studies

The 3D structure of different OsFD7 protein was predicted utilizing Robetta server (http://robetta.bakerlab.org/; Kim *et al*., 2004). Best 3D model was selected by performing Ramachandran plot analysis available at MOLProbity server (http://molprobity.biochem.duke.edu/). These structures were visualized with the help of Jmol tool (http://www.jmol.org/). STRING database 10.0 (Szklarczyk *et al*., 2015) was utilized for identifying putative interacting partners of OsFD7. PSORT prediction software was used for predicting the cell compartment where OsFD7 is localized (http://psort1.hgc.jp/form.html).

All the assays/experiments performed in this study were repeated at least thrice. All primer sequences are given in Table S1 and S2.

## RESULTS

### Expression profiling of *OsFD7* at different stages of rice panicle initiation and development

The microarray analysis revealed that *OsbZIP62*/*OsFD7* expresses preferentially in SAM and during very early stages of panicle development, i.e. P1-1, P1-2 and P1-3. Its transcript declined gradually from P1 to P6 stages of panicle development (Fig. 1A); P1-1, P1-2 and P1-3 form a sub-set of P1 (see Nijhawan *et al*., 2008; Sharma *et al*., 2012); *OsFD7* transcript level was much lower in vegetative tissues. This microarray profile could be validated by real time PCR and fairly matched with western blot analysis (Fig. 1B,C). An additional high molecular weight band was visible specifically in the SAM protein sample in the western blot (Fig. 1C) that could represent the phosphorylated form of OsFD7. Additionally, we compared the expression profile of *OsFD7* with all previously identified *OsFDs* (Taoka *et al*., 2011; Tsuji *et al*., 2013; Cerise *et al*., 2020). *OsFD1*, *OsFD2* and *OsFD4* genes express at very low levels, however, *OsFD3* and *OsFD6* genes express at high levels at almost all the stages of development analyzed with slight variability in the expression pattern (Figure 1A; Figure S12); *OsFD5* was not represented in the Affymetrix microarray chip used then by our group (Nijhawan *et al*., 2008).

**Figure 1.**
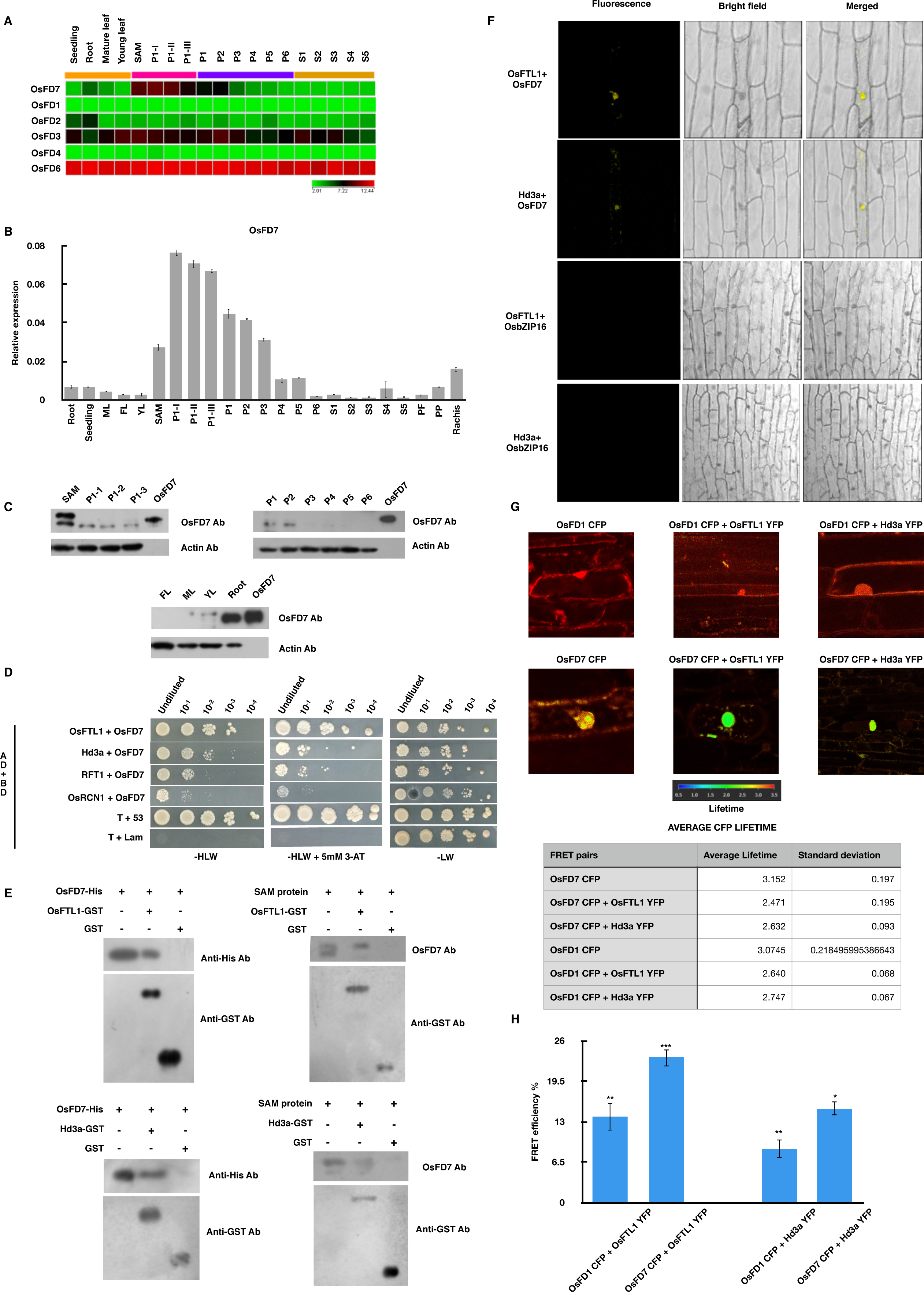
Expression profile of *OsFD7* during different stages of rice panicle development and its interaction with OsFTLs and OsRCN1. A: Heat maps depicting expression profile of *OsFD7* and *OsFDs* during different stages of rice development (GSE6893) as analyzed by microarray (retrieved from Nijhawan et al., 2008). **B:** Real-time PCR analysis of *OsFD7* in the tissue samples used in **A** above. *UBQ5* and *eEF-1* genes were used as internal controls (Jain et al., 2006). Error bars represent standard error. (Refer experimental procedures for detailed information about different developmental stages). **C:** Protein profile of *OsFD7* at different developmental stages. Western blot for vegetative tissues (used as control) probed with OsFD7 antibody was overexposed. Actin was used as a loading control for all the western blots. OsFD7 lane corresponds to bacterially expressed OsFD7 protein with T7 and His tag. **D:** Y2H assay - Positive interaction observed between OsFD7 and OsFTLs and OsRCN1. Positive control: pGADT7-T + pGBKT7-53; Negative control: pGADT7-T + pGBKT7-Lam. AD and BD denotes activation domain and DNA binding domain of GAL4, respectively. -HLW+3-AT: selective medium (SD/-His-Leu-Trp) supplemented with 3-AT; -LW: control medium (SD/-Leu-Trp). **E:** GST pull-down assay confirming interaction of OsFD7 with OsFTLs. OsFD7-His: His tagged protein; OsFTL1-GST, Hd3a-GST: GST tagged OsFTL1 and Hd3a protein; GST: GST protein. **F:** BiFC assay showing strong interaction between OsFD7 and OsFTLs and the formation of a stable hetero-dimer inside nucleus. Negative controls: OsFTL1 + OsbZIP16 and Hd3a + OsbZIP16. **G:** FAST-FLIM images of different constructs after particle bombardment in onion peel cells depicting CFP lifetime of the different protein combinations with a common colored scalebar. The fluorescence lifetime is depicted by a false colour representation. Average CFP lifetime for different FRET pairs is tabulated in the form of a table. **H:** Bar graphs representing FRET efficiencies of different samples. Error bars represent standard error (*: *p* < 0.05, **: *p* < 0.01, ***: *p* < 0.001, Student’s *t*-test).

### Sequence analysis and phylogenetic relationship of OsFD7 with FD and FD-like proteins

The domain analysis using SMART software confirmed the presence of a typical bZIP domain (179 to 248 aa) in OsFD7 protein (Fig. S1B); structural analysis, localization and homodimerization potential of OsFD7 are described in detail in appendix S2 of supporting information. Further, SALAD database predicted the occurrence of five additional motifs in OsFD7 conserved across different FD proteins (Fig. S4A). Among these, motif 3 is present in almost all FDs identified from different plant species and contains a potential CDPKs phosphorylation site (L-X-R/K-X-X-S/T). Moreover, OsFD7 contains motif 4 and 7, which are well conserved in AtFD, SlFDs and some TaFDs but, surprisingly, motif 7 is not present in other FDs identified from rice (Fig. S4A).

The protein sequences of FD and FD-like genes retrieved from phytozome were aligned using MAAFT server to identify the closest ortholog of OsFD7. Highest conservation was observed in the region harbouring bZIP domain and motif 3 at the C-terminal end, with slight differences in the aa sequences of these proteins (Fig. S4B). The OsFD7 protein contains non-canonical SAS aa sequence in motif 3 instead of canonical SAP/TAP aa present in OsFD1/AtFD proteins that is recognized by GF14-3-3 protein (Fig. S4B) (Abe *et al*., 2005; Taoka *et al*., 2011). The phylogenetic tree constructed using PhyML 3.0 server, for FD gene sequences of different plant species retrieved using phytozome v12.1 (Table S3), clearly showed that OsFD7 aligns with the monocot clade with highest homology with TaFDL2, TaFDL3 and TaFDL6 of *Triticum aestivum* (Fig. S4C).

### Interaction of OsFD7 with OsFTL proteins

To examine if OsFD7, like other FD or FD-like protein in rice, interacts with OsFTLs, among the 13 FT-like protein coding genes identified in rice, three FTLs were selected for this study: OsFTL1, Hd3a (OsFTL2) and RFT1 (OsFTL3). Additionally, we checked its interaction with OsRCN1, a closest ortholog of AtTFL1 (Nakagawa *et al*., 2002). Y2H analysis revealed that OsFD7 interacts more strongly with OsFTL1 followed by Hd3a and RFT1; however, a very weak interaction was observed with OsRCN1 (Fig. 1D). The strongest interaction of OsFD7 with OsFTL1 is not surprising because of their overlapping expression domains (Fig. S5A). Vector swapping experiments could not be performed as Hd3a and RFT1 exhibit strong transactivation potential (Fig. S5B). These interactions were verified by GST pull-down assays (Fig. 1E). In addition, SAM protein extracts were used to perform *in vitro* GST pull-down assay. Strikingly, OsFTL1 and Hd3a both interacted preferentially with the upper band of OsFD7 protein, possibly representing its phosphorylated form (Fig. 1E). Subcellular localization and BiFC experiments were performed in onion peel cells using biolistic particle delivery system. Although OsFTL1 and Hd3a proteins localize to both cytoplasm as well as the nucleus (Fig. S5C) but their interaction with OsFD7 was mainly detected in the nucleus where OsFD7 predominantly localizes (Fig. 1F, Fig. S2C). Also, FLIM-FRET analysis was carried out to measure the CFP lifetime of different FRET pairs as well as to determine their FRET efficiencies. OsFD7-OsFTL1 (23.29%) and OsFD7-Hd3a (15.1%) protein pairs yielded a higher FRET efficiency than OsFD1-OsFTL1 (13.7%) and OsFD1-Hd3a protein pairs (8.7%) (Fig. 1G,H).

### Interaction of OsFD7 with OsGF14 protein

To examine if OsFD7 interacts with OsGF14-3-3 protein, Y2H assay was performed and it revealed strong interaction with OsGF14b,c,d proteins (Fig. S7A). Moreover, the near constitutive expression of OsGF14b,c,d (Fig. S6A) at SAM and during panicle development as well as the presence of GF14-3-3 binding site at the C-terminal end of OsFD7 protein (R/K-X-X-S-X-S) further indicated the possibility of their interaction, although this site is slightly different from the canonical one (R/K-S-X-P and R/K-X-X-S-X-P) (Fig. S4B). The interactions between OsGF14b,c,d and OsFTL1, Hd3a, RFT1 were also confirmed by both Y2H (Fig. S7A) and GST pull-down assays. As clearly visible in the western blot, OsGF14b interacted preferentially with the probable phosphorylated form of OsFD7 (Fig. S7B). The BiFC assay carried out in onion peel cells supported the above observations. Notably, BiFC signal for OsFD7-OsGF14b interaction was detected in the nucleus whereas OsGF14b protein alone is localized predominantly in the cytoplasm with weak signal in the nucleus. In comparison, BiFC signal for OsGF14b-OsFTL1 interaction was visible in the cytoplasm where OsGF14b is predominantly detected (Fig. S7C, Fig. S6B).

To delineate the region of OsFD7 that interacts with OsGF14s, ΔbZIP and ΔC-ter constructs of OsFD7 generated were employed for Y2H analysis (Fig. 2A). The ΔbZIP construct could interact whereas ΔC-ter construct failed to interact with OsGF14s (Fig. 2B). The GST pull-down assay provided essentially similar results (Fig. 2C). To further substantiate these results, 84 bp C-terminal region (named as C-pep WT) amplified from OsFD7 was used to carry out Y2H and GST pull-down assays with OsGF14b and a positive interaction could be observed (Fig. 2A,B,C) suggesting that this C-terminal peptide indeed interacts with OsGF14b protein and is sufficient for this interaction to occur.

**Figure 2.**
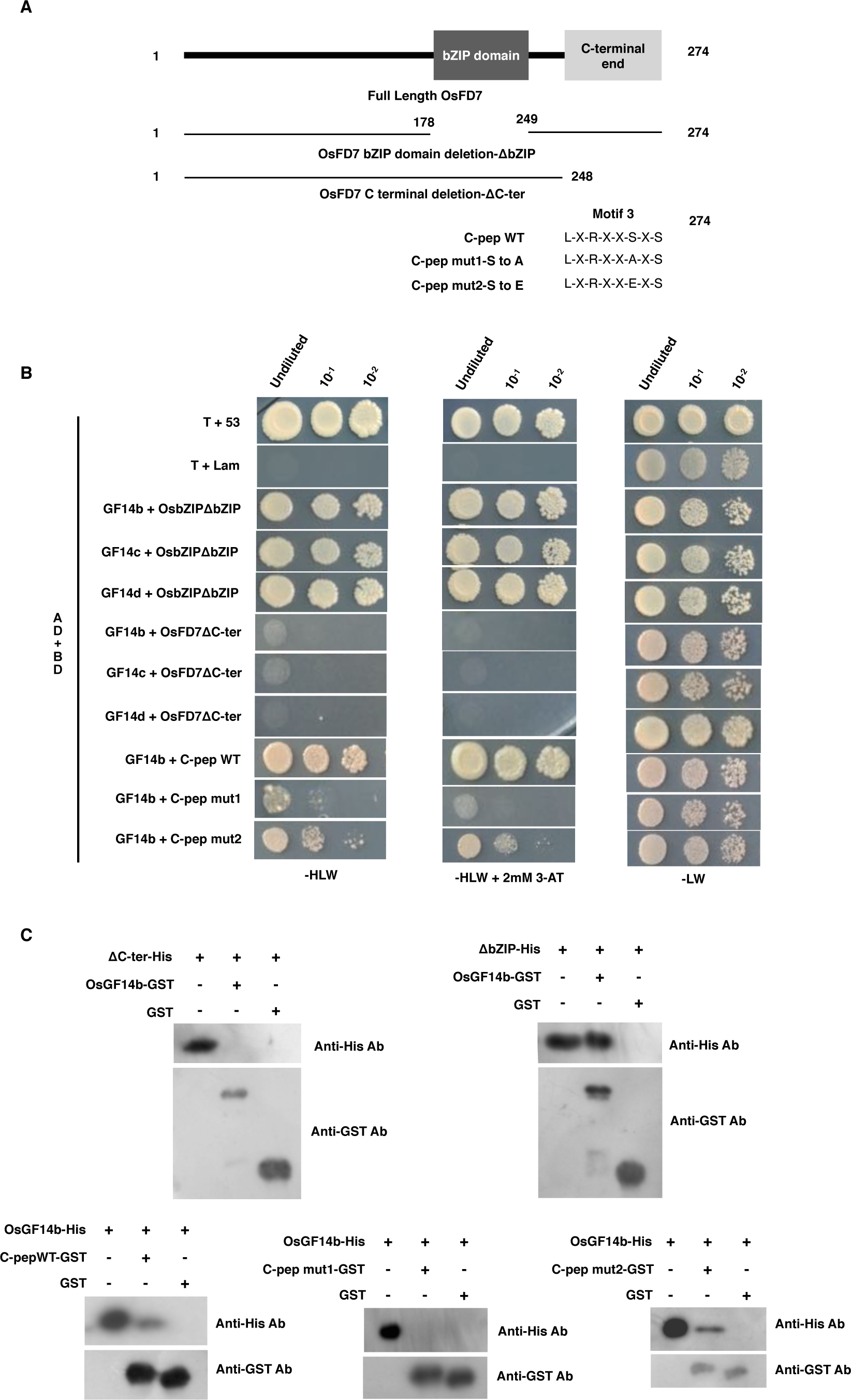
Interaction of OsGF14s with truncated protein and mutated peptides of OsFD7. **A:** Diagrammatic representation of truncated OsFD7 protein as well as mutated peptide of OsFD7 harbouring amino acid substitutions. **B:** Y2H assay using truncated OsFD7 and mutated OsFD7 peptides. Positive control: pGADT7-T + pGBKT7-53; Negative control: pGADT7-T + pGBKT7-Lam. AD and BD denotes activation domain and DNA binding domain of GAL4, respectively. -HLW+3-AT: selective medium (SD/-His-Leu-Trp) supplemented with 3-AT; - LW: control medium (SD/-Leu-Trp). **C:** GST pull-down assay performed with truncated OsFD7 bZIP-His: bZIP domain deleted OsFD7 His tagged protein; ΔHis: C-terminal truncated OsFD7 His tagged protein; C-pep WT,mut1,mut2-GST: GST tagged mutated OsFD7 C terminal peptides; GST: GST protein.

To evaluate whether the interaction of OsGF14b and OsFD7 is phosphorylation dependent, as C-terminal end of OsFD7 harbours a potential CDPK phosphorylation site (L-X-R/K-X-X-S) overlapping with the non-canonical GF14-3-3 binding site (R/K-X-X-S-X-S) (Fig. S4B), two mutant constructs containing 84 bp C-terminal region were amplified from full-length OsFD7. One of the mutants (named as C-pep m1-StoA) contains a non-phosphorylatable substitution of serine with alanine while the other one (named as C-pep m2-StoE) contains a phospho-mimic substitution of serine with glutamate (Fig. 2A). In the Y2H assay performed with OsGF14b. C-pep m1 failed to interact with OsGF14b while C-pep m2 displayed positive interaction (Fig. 2B). Similar results were obtained with GST pull-down assay too (Fig. 2C).

### Formation of a tri-protein complex involving OsFD7, OsGF14 and OsFTL proteins

From the results presented in the previous sections, it could be presumed that OsFD7, OsGF14s and OsFTLs proteins might be interacting with each other to form a ternary complex *in vivo*. To examine if such an interaction occurs, yeast-3-hybrid system (Y3H) was employed. The Y3H approach makes use of a pBRIDGE vector harbouring two MCS; *OsFD7* gene was cloned in MCSI while *OsGF14b,c* genes were cloned separately in MCSII, to determine whether OsGF14b,c bridges interaction between OsFTLs and OsFD7. These pBRIDGE vector constructs were co-transformed with pGADT7 vector constructs, containing cloned *OsFTL* genes, in AH109 yeast strain. Strong interactions were observed in all the cases (Fig. S7D).

### Direct interaction of OsFTL proteins with OsFD7

To examine whether the interaction between OsFD7 and OsFTLs is mediated by OsGF14 protein(s), we performed Y2H assay using ΔC-ter construct of OsFD7 as OsGF14 proteins recognize serine residue at the C-terminal end of OsFD7; its deletion would otherwise abolish the interaction between OsFTLs and OsFD7 if it is mediated by endogenous OsGF14s. To our surprise, a direct interaction was observed with both OsFTL1 and Hd3a (Fig. 3A); however, OsFTL1 interacted more strongly with OsFD7 in comparison to Hd3a in Y2H assay (Fig. 3A). These results could be validated by GST pull-down assay (Fig. 3B). To further substantiate these results, mutant C-terminal constructs of OsFD7, StoA (mut1) and StoE (mut2) (Fig. 3C), were used for Y2H analysis. Strong interactions were observed between OsFTL1 and mutated versions of OsFD7, however, a weak interaction could be seen with Hd3a (Fig. 3D); see appendix S2 for result of additional experiment supporting the direct interaction between OsFTLs and OsFD7.

**Figure 3.**
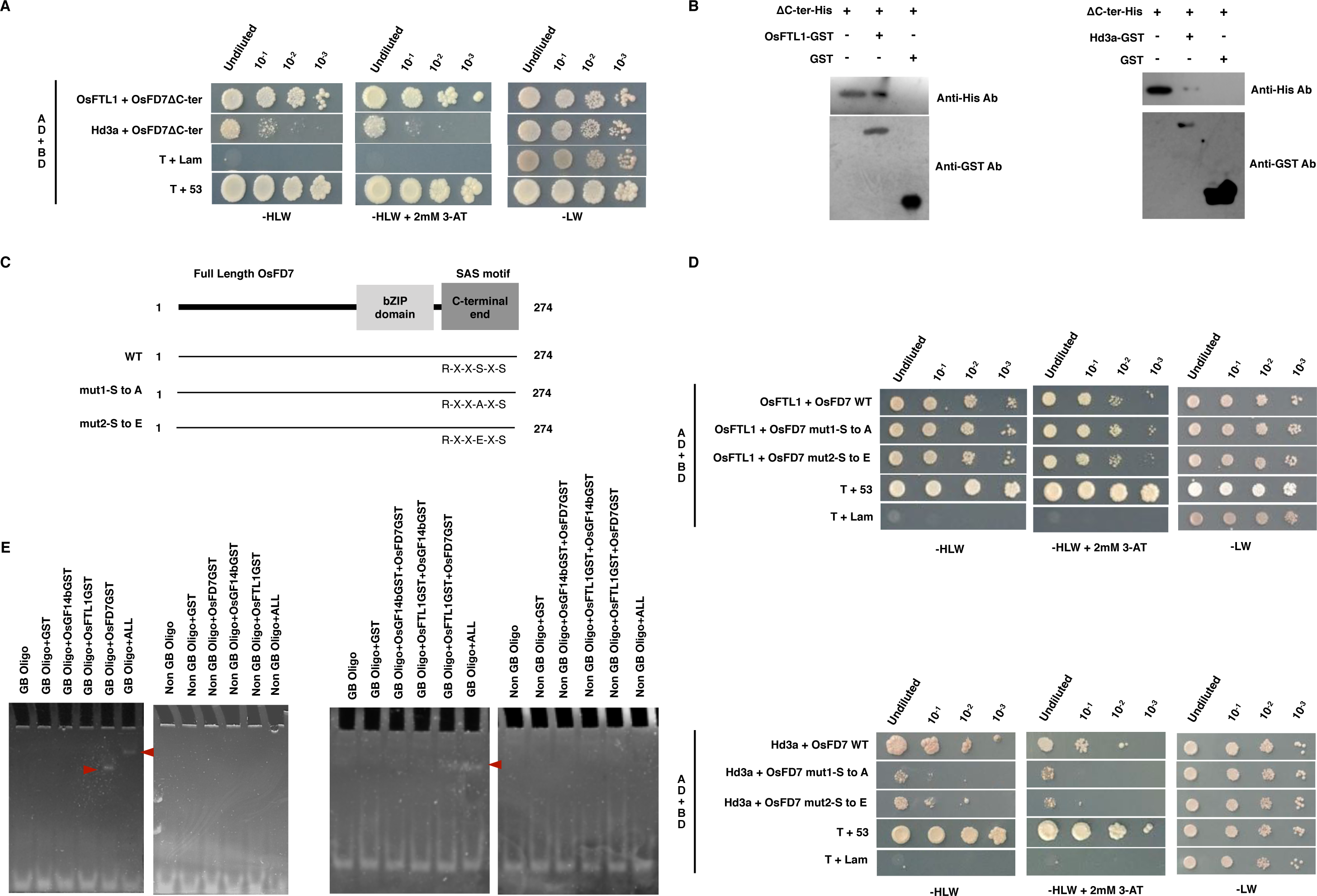
Interaction of OsFTLs with full length and truncated/mutated OsFD7. **A:** Y2H assay depicting protein-protein interactions between OsFTLs and truncated OsFD7 protein. Positive control: pGADT7-T + pGBKT7-53; Negative control: pGADT7-T + pGBKT7-Lam. AD and BD denotes activation domain and DNA binding domain of GAL4, respectively. - HLW+3-AT: selective medium (SD/-His-Leu-Trp) supplemented with 3-AT; -LW: control medium (SD/-Leu-Trp). **B:** GST pull-down assay with truncated OsFD7. ΔOsFD7 His tagged protein without C-terminal region; OsFTL1-GST: GST tagged OsFTL1 protein; Hd3a-GST: GST tagged Hd3a protein; GST: GST protein. **C:** Diagrammatic representation of OsFD7 mutated protein harbouring amino acid substitutions. **D:** Y2H assay showing interactions between OsFTLs and mutated OsFD7 proteins. Positive control: pGADT7-T + pGBKT7-53; Negative control: pGADT7-T + pGBKT7-Lam. AD and BD denotes activation domain and DNA binding domain of GAL4, respectively. -HLW+3-AT: selective medium (SD/-His-Leu-Trp) supplemented with 3-AT; -LW: control medium (SD/-Leu-Trp). **E:** DNA mobility shift assay to determine interaction of G-box element with different components. OsFD7GST: GST tagged OsFD7 protein; OsGF14bGST: GST tagged OsGF14b protein; OsFTL1GST: GST tagged OsFTL1 protein; ALL: OsFD7GST + OsFTL1GST + OsGF14bGST; GST: GST protein. GB oligo: G-box element oligo; Non GB oligo: Non G-box element oligo. DNA mobility shifts marked with red arrows.

### DNA binding activity of OsFD7

To confirm DNA binding ability of OsFD7, mobility shift assay was performed with G-box and C-box elements. Two oligonucleotides were selected: one with G-box (GB oligo) and another without G-box (Non-GB oligo). A shift in the band could be detected in the lane containing GB oligo+OsFD7GST confirming that OsFD7 binds G-box element (Fig. 3E). However, no shift could be seen in in the lanes containing GB Oligo+OsGF14bGST and GB Oligo+OsFTL1GST. We observed a super shift in the band upon addition of all three proteins to GB oligo (Fig. 3E). To identify the combination of proteins mainly responsible for this super shift, different protein pairs were tested in the gel-shift assay. Protein pair OsFTL1GST+OsFD7GST super shifted GB Oligo; however, lanes containing other protein combinations showed no further shift in the band (Fig. 3E). No bands could be seen in Non-GB oligo gels. Essentially similar results were obtained when mobility shift assays were performed with CB oligos (harbouring C-box elements) and Non-CB oligos (Fig. S8B).

### OsCDPK41 and OsCDPK49 interact with and phosphorylate OsFD7

*In silico* predictions using STRING database identified some additional proteins like OsCDPKs that might be interacting with OsFD7 (Fig. S9A). Among these, *OsCDPK41* and *OsCDPK49* were selected for further characterization expressing at almost all the developmental stages of rice analysed by microarray (Fig. S9B). CDSs of *OsCDPK41* and *49* were cloned in yeast expression vectors and Y2H assay performed clearly showed that they do interact with OsFD7 (Fig. 4A). As OsCDPK49 exhibited strong transactivation activity, vector-swapping experiments could not be performed (Fig. S9C). These interactions could also be observed in GST pull-down assays and that too without any requirement for calcium (Fig. 4B). In the BiFC assay performed using onion peel cells, the signal for OsFD7-OsCDPK41 and OsFD7-OsCDPK49 interactions could be detected inside the nucleus (Fig. 4C) where OsFD7 is mainly localized (Fig. S2C), whereas OsCDPK41 and OsCDPK49 localize both in the nucleus as well as cytoplasm (Fig. S9D).

**Figure 4.**
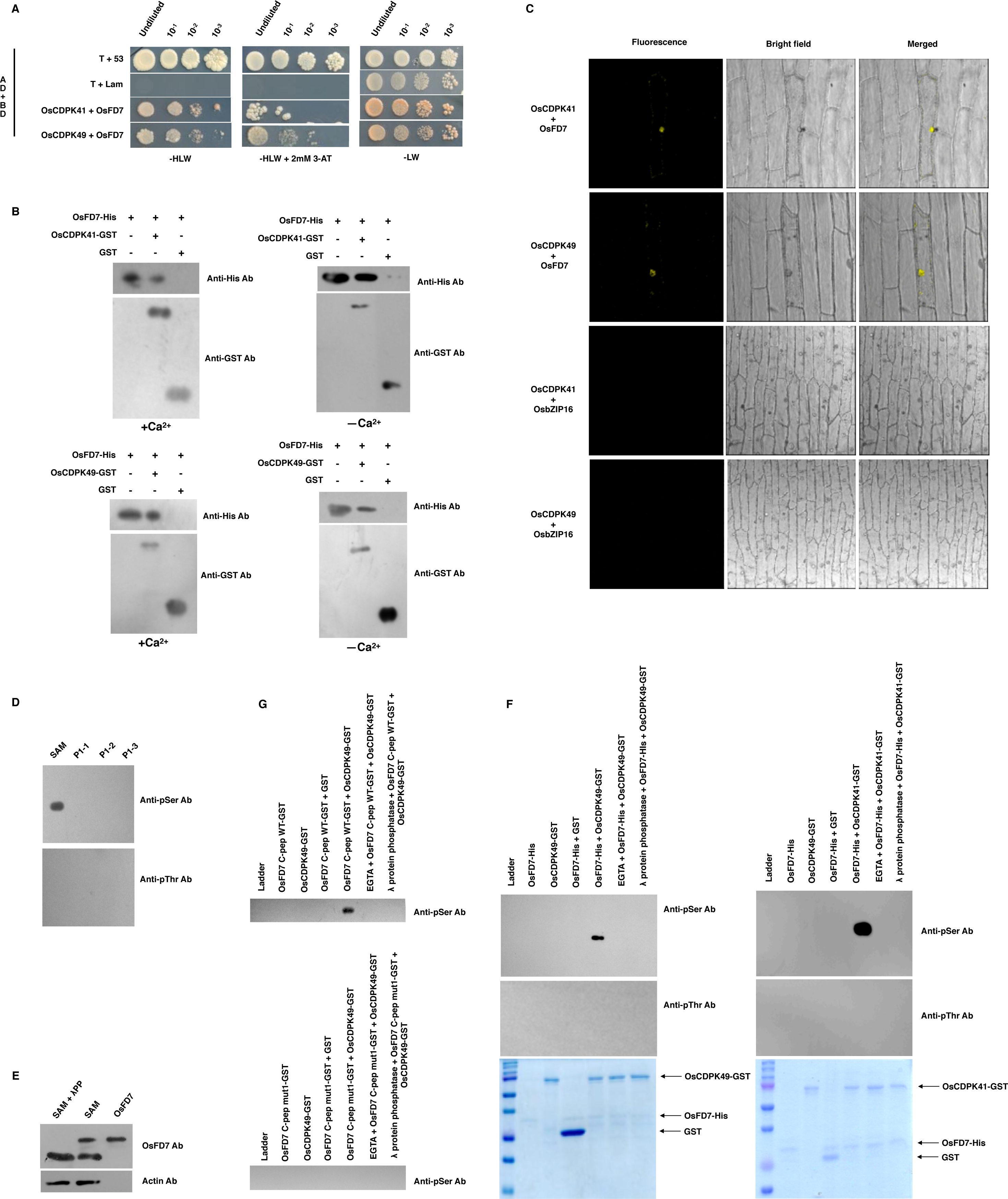
Interaction of OsFD7 with OsCDPKs. **A:** Y2H assay-Positive interaction observed in between OsFD7 and OsCDPKs. Positive control: pGADT7-T + pGBKT7-53, Negative control: pGADT7-T + pGBKT7-Lam. AD and BD denotes activation domain and DNA binding domain of GAL4, respectively. -HLW+3-AT: selective medium (SD/-His-Leu-Trp) supplemented with 3-AT, -LW: control medium (SD/-Leu-Trp). **B:** GST pull-down assay confirming interaction of OsFD7 with OsCDPKs in the presence or absence of calcium. OsFD7-His: His tagged OsFD7 protein, OsCDPK41-GST: GST tagged OsCDPK41 protein, OsCDPK49-GST: GST tagged OsCDPK49 protein, GST: GST protein. **C:** BiFC assay showing interaction between OsFD7 and OsCDPKs. BiFC signal is detected in the nucleus. Negative controls: OsCDPK41+OsbZIP16 and OsCDPK49+OsbZIP16. **D:** *In vivo* phosphorylation detection assay by immunoprecipitation with OsFD7 antibody. Strong signal detected in SAM tissue with anti-phosphoserine antibody using OsFD7 immuprecipitated samples. No band was obtained with anti-phosphothreonine antibody. **E:** Protein dephosphorylation assay using λ protein phosphatase. OsFD7 protein was immmunoprecipitated from SAM tissue using OsFD7 antibody and treated with λ protein phosphatase. Western blot was probed with OsFD7 antibody. Actin was used as a loading control. OsFD7 lane corresponds to bacterially expressed OsFD7 protein with T7 and His tag **F:** *In vitro* kinase assay showing phosphorylation of OsFD7 by OsCDPK49 and OsCDPK41 using anti-phosphoserine/phosphothreonine antibody. A strong signal could be seen in 5th lane with anti-phosphoserine antibody. OsFD7-His: His tagged OsFD7 protein; OsCDPK49,41-GST: GST tagged OsCDPK49,41 protein; GST: GST protein. **G:** *In vitro* kinase assay showing phosphorylation of GST tagged C-terminal peptide of OsFD7 by OsCDPK49 using anti-phosphoserine. A strong signal could be seen in 5th lane with anti-phosphoserine antibody with C-pep WT-GST while no band could be seen wherein mutated versions of C-terminal peptide of OsFD7 was used for assay. OsFD7 C-pep WT-GST: OsFD7 C-terminal GST tagged WT peptide; OsFD7 C-pep mut1-GST: OsFD7 C-terminal GST tagged peptide with a single amino acid substitution StoA; OsCDPK49-GST: GST tagged OsCDPK49 protein; GST: GST protein.

The presence of a potential CDPK phosphorylation site (L-X-R/K-X-X-S) at its C-terminal end (Fig. S4B) and extra band observed in the lane of SAM protein (Fig. 1C), provided clue that it most likely represents the phosphorylated form of OsFD7. Thus, *in vivo* phosphorylation status of OsFD7 protein was examined. Protein extracts of SAM, P1-1, P1-2 and P1-3 tissues were immunoprecipitated with anti-OsFD7 antibody followed by western blot analysis; only anti-phosphoserine antibody probed blot showed the presence of a sharp band in the SAM lane (Fig. 4D). Moreover, the upper band visible in Fig. 1C disappeared when OsFD7 protein immunoprecipitated from SAM was treated with lambda protein phosphatase confirming that it indeed represents the phosphorylated form of OsFD7 (Fig. 4E). Also, an *in vitro* kinase assay was performed using OsCDPK49 and OsCDPK41 purified protein against the purified full-length OsFD7 protein acting as a substrate. The reaction mixtures with appropriate controls, including autophosphorylation controls, were subjected to western analysis. Phosphorylated OsFD7 bands could be detected in the presence of calcium with anti-phosphoserine antibody but not with anti-phosphothreonine antibody. The addition of EGTA to the kinase reaction mixture abolished the phosphorylation of OsFD7 by these CDPKs, indicating that it is calcium-dependent. Also, addition of Lambda protein phosphatase to the reaction mixture reversed the phosphorylation of OsFD7 (Fig. 4F).

To confirm the potential CDPK phosphorylation site, we performed an *in vitro* kinase assay using OsCDPK49 purified protein in the presence of calcium with the purified WT and mutated version of C-terminal GST tagged peptides of OsFD7. A signal could be detected after western blotting in the 5^th^ lane containing the C-terminal GST tagged WT peptide and not with C-terminal GST tagged mut1 (StoA substitution) peptide of OsFD7 (Fig. 4G); appropriate controls were included in the experiment. The supporting information to dissect out which region of OsFD7 is responsible for its interaction with OsCDPKs as well as to confirm the requirement of leucine at -5 position in L-R-R-T-T-S sequence at C-terminal end of OsFD7 is provided in appendix S2.

### *OsFD7* RNAi transgenics are late flowering and exhibit altered panicle architecture

To gain insight into the functional significance of OsFD7 in its native plant species, RNAi transgenics were raised in PB-1 variety of rice (*Oryza sativa indica*). Out of six homozygous lines, three were selected for detailed functional characterization. All three RNAi lines had reduced levels of OsFD7 (Fig. 5A,B) and flowered later than vector control (VC) plants (transformed with empty pANDA vector) (Fig. 5C,D).

**Figure 5.**
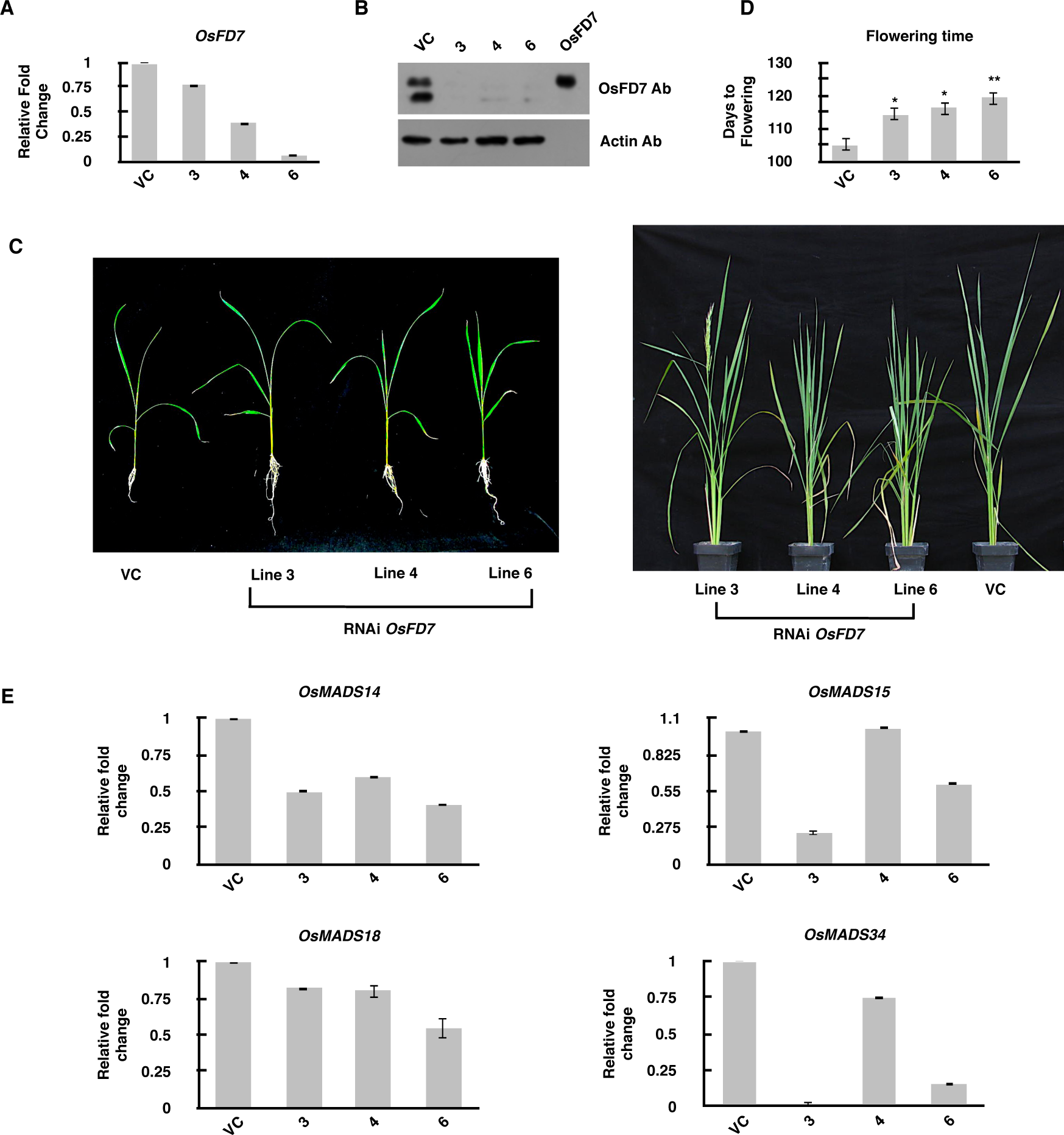
Characterization of *OsFD7* RNAi rice transgenics. **A:** Real-time PCR analysis of *OsFD7* in different transgenic lines with respect to VC at SAM. *UBQ5* and *eEF-1*α genes were used as internal controls (Jain et al., 2006). Error bars represent standard error. **B:** Western blot showing protein levels of OsFD7 in different transgenic lines and VC plants. Actin was used as a loading control. OsFD7 lane corresponds to bacterially expressed OsFD7 protein with T7 and His tag. **C:** Phenotypic characterization of *OsFD7* RNAi transgenics at different stages of plant development. **D:** Graphical representation of flowering times of transgenic plants with respect to VC plants (mean ± SD) (*: *p* < 0.01, **: *p* < 0.001, Student’s *t*-test). **E:** Real-time PCR analysis of known flowering genes in different transgenic lines with respect to VC at SAM. *UBQ5* and *eEF-1* genes were used as internal controls (Jain et al., 2006). Error bars represent standard error.

To identify the regulatory targets of OsFD7, we isolated RNA from SAM tissue individually from these RNAi transgenics and VC plants and performed real-time PCR for some known floral meristem identity genes. There was significant down-regulation in the expression levels of *OsMADS14*, *OsMADS15* and *OsMADS18* genes, which belong to *AP1/FUL*-like subfamily of MADS box genes (Preston and Kellogg, 2006; Kobayashi *et al*., 2012), in almost all the lines tested (Fig. 5E). The transcript level of *Panicle Phytomer2* (*OsPAP2/OsMADS34*) belonging to *Sepallata* (*SEP*) gene subfamily of MADS box transcription factors, also decreased considerably in RNAi transgenics (Fig. 5E). The details about the specificity of RNAi construct are provided in appendix S2.

Besides late flowering, *OsFD7* RNAi transgenics of rice displayed altered inflorescence morphology, with slightly longer and dense panicles with more florets in comparison to VC plants (Fig. 6A,B,C,D). The transcript abundance of *OsMADS34*, which has an influence on the number of primary and secondary branches as well as number of spikelets (Gao *et al*., 2010; Kobayashi *et al*., 2012), was significantly down-regulated in SAM and early panicle stages of these RNAi lines (Fig. 5E, Fig. 7). Although, size and weight of the seeds was slightly higher in these transgenics (Fig. 6E,F), however, the number of unfilled seeds was more in these RNAi lines vis-à-vis VC plants (Fig. 6D). The transcript levels of the members of *SEP* gene family, including *OsMADS34*, *OsMADS1*, *OsMADS5*, *OsMADS7* and *OsMADS8*, that are known to control branch, spikelet and floral meristem development and floral organ initiation and differentiation (Cui *et al*., 2010; Gao *et al*., 2010), were also quantitated at the early stages of panicle development. Except *OsMADS5*, the transcript levels of *OsMADS1*, *OsMADS7, OsMADS8* and *OsMADS34* were reduced (Fig. 7).

**Figure 6.**
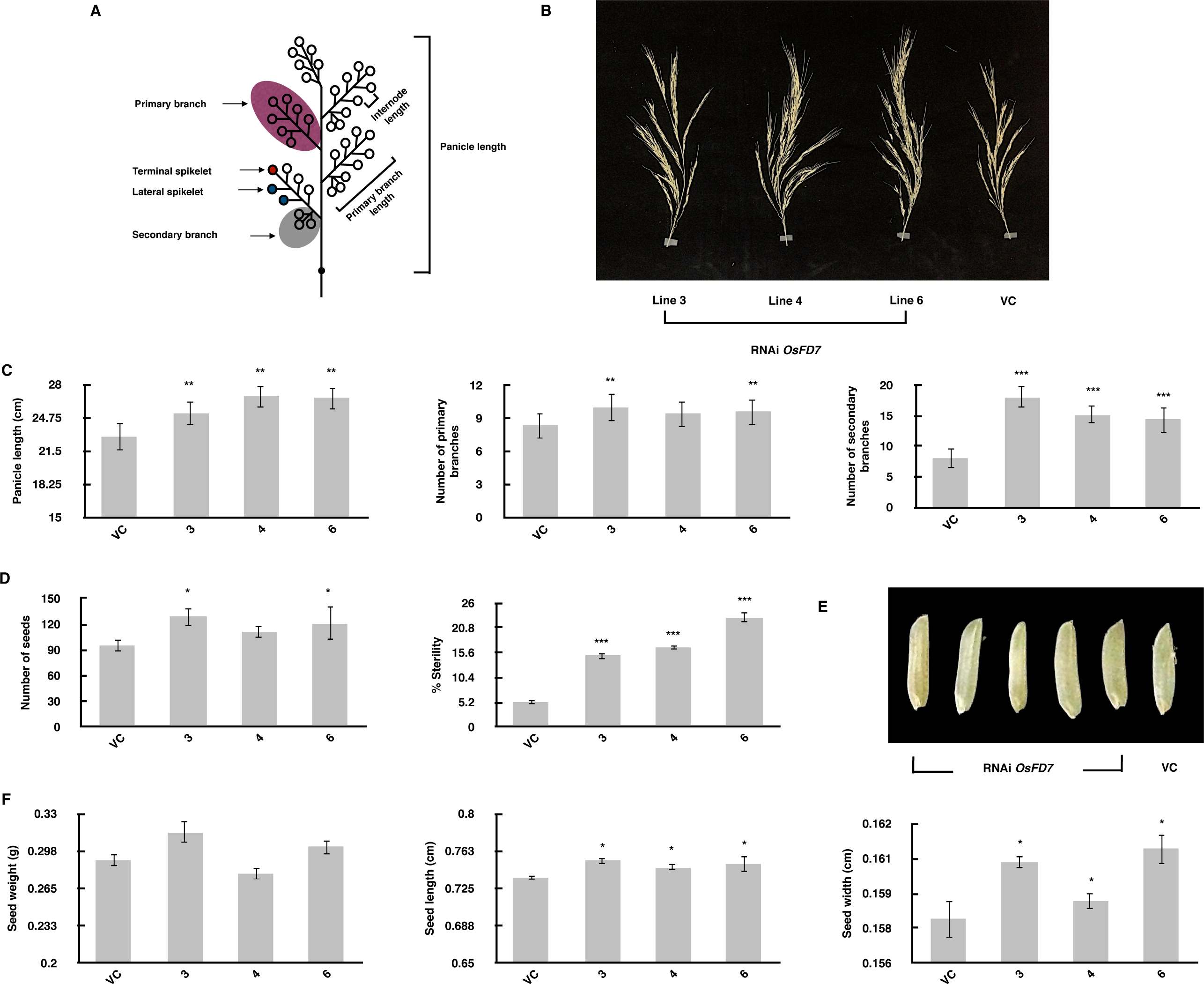
Changes in panicle morphology of *OsFD7* RNAi rice transgenics. **A:** Diagrammatic representation of rice panicle. **B:** Difference in the panicle length of *OsFD7* RNAi transgenics in comparison to VC plants. **C:** Graphical representation of differences in the panicle length, number of primary and secondary branches in transgenics. **D:** Graphical representation of differences in the seed number and percent sterility in transgenics plants. **E:** Seeds of transgenics and VC plants. **F:** Graphical representation of difference in seed weight and seed dimensions of transgenics. (mean ± SD) (*: *p* < 0.05, **: *p* < 0.01, ***: *p* < 0.001, Student’s *t*-test).

**Figure 7.**
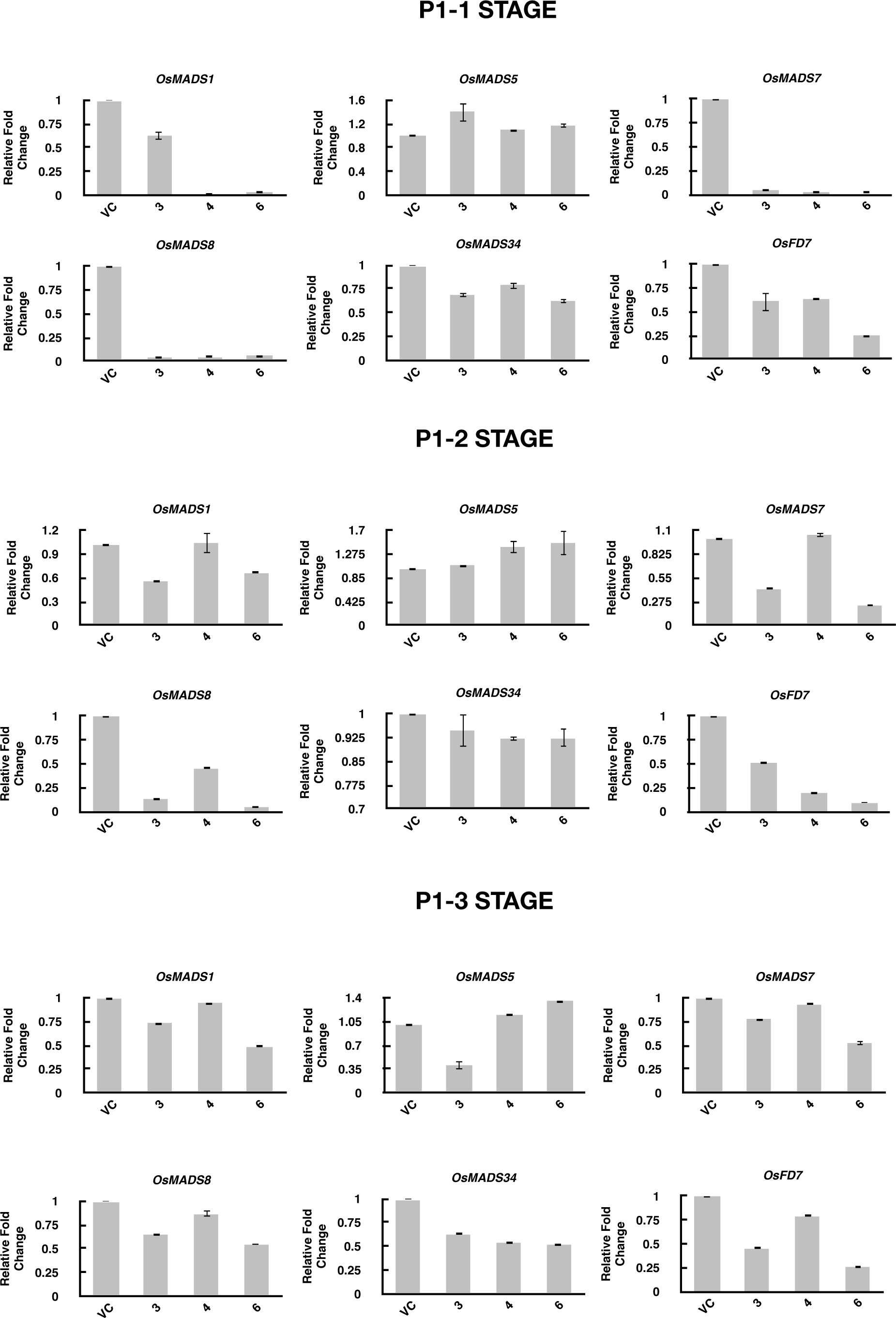
Real-time PCR analysis of *OsFD7* and known floral meristem and organ identity genes in *OsFD7* RNAi transgenic and VC plants at **(A)** P1-1, **(B)** P1-2, **(C)** P1-3 stages of panicle development in rice. *UBQ5* and *eEF-1* genes were used as internal controls (Jain et al., 2006). Error bars represent standard error.

## DISCUSSION

### OsFD7 expresses predominantly in SAM and during early stages of panicle development

The microarray-based transcriptome analysis revealed that the expression of *OsbZIP62*/*OsFD7* is considerably high in SAM and during early stages of panicle development; it has virtually little expression in the vegetative tissues examined. In comparison, *OsFD1*, *OsFD2* and *OsFD4* express at low level, more or less uniformly, whereas *OsFD3* and *OsFD6* express at higher levels in all tissues examined (Fig 1A; see also Nijhawan *et al*., 2008). The presence of relatively higher levels of OsFD7 protein in tissue samples representing SAM and early panicle stages in rice almost matches with its transcript profile and this makes us speculate that it may play a predominant role in rice reproductive development.

### Similarity of OsFD7 with FD and FD-like proteins

In addition to the presence of a bZIP domain, OsFD7 showed the existence of five additional motifs well spread throughout the protein. Apart from motif 3 that occurs in almost all the previously identified FDs from both monocots and dicots involved in flowering pathway, OsFD7 additionally harbours motifs 4 and 7, which are well conserved in the FD proteins of *Arabidopsis thaliana*, *Solanum lycopersicum* and *Triticum aestivum*, clearly indicating its close proximity to AtFD, SlFDs and TaFDs. Surprisingly, other rice FDs lack motif 7. In fact, OsFD7 contains an extra motif 6 when compared to AtFD. Additionally, OsFD7 showed highest similarity with TaFDLs in comparison to other monocot FDs.

In depth sequence analysis of these motifs showed the presence of threonine aa specifically in motif 3 (TXX) of dicots in comparison to serine aa (SXX) in monocots. An earlier study on OsFDs have claimed that slight changes observed in the aa sequence of the motifs shared by these FDs was largely responsible for the functional divergence of OsFD2 from OsFD1 (Tsuji *et al*., 2013). Although OsFD7 shared significant similarity with these FDs, but there were slight differences in the aa sequence of these conserved motifs; for example, motif 3 of OsFD7 contains SAS aa instead of SAP/TAP/TAP aa present in OsFD1/AtFD/SlFD sequence; in fact, TaFDs harbour SAP/STS aa sequence in motif 3. The presence of this non-canonical sequence within the GF14-3-3 binding site and the presence of additional motifs could possibly contribute to the neo-functionalization of OsFD7 in comparison to AtFDs and other OsFDs identified till date, although this assumption requires experimental validation.

### Interaction of OsFD7 with the components of FAC

An earlier study by Taoka *et al*. (2011) showed that the bZIP protein OsFD1 interacts with Hd3a *via* GF14-3-3 protein to form a functional FAC at SAM initiating transition from vegetative to reproductive development. In this study, OsFD7 did indeed show interaction with OsFTLs; the interaction was strongest with OsFTL1, followed by Hd3a and RFT1. This was further confirmed by FLIM-FRET analysis wherein OsFD7-OsFTLs exhibited higher FRET efficiency in comparison to OsFD1-OsFTLs. Additionally, both OsFTL1 and Hd3a interacted preferably with the phosphorylated form of OsFD7. The strong interaction of OsFD7 with OsFTL1 and their overlapping expression profiles suggest that they might be performing some specialized functions at SAM and during early stages of panicle development, apart from their probable role in floral evocation pathway. In fact, a recent report by Brambilla *et al*. (2017) also showed interaction between OsFD7 and Hd3a, without any further elaboration.

Considering the high expression OsFD7 has in SAM, one could extrapolate that these interactions most likely take place in the nucleus of the cells of shoot apex, as BiFC signal was mainly detected in the nucleus of onion peel cells. Moreover, a weak interaction observed between OsFD7 and OsRCN1 (homologous to AtTFL1) suggests that OsFD7 along with OsRCN1 might repress flowering in rice similar to the case observed in *Arabidopsis* wherein binding of FD with FT activates flowering while its association with TFL1 represses flowering depending on FT/TFL1 ratio (Ahn *et al*., 2006; Wickland and Hanzawa, 2015; Moraes *et al*., 2019).

Further, OsFD7 showed positive interaction with other functional components of FAC, i.e. OsGF14s, in the nuclei of onion peel cells, indicating that OsGF14 protein is recruited to the nucleus by OsFD7; OsGF14 alone is weakly localized in the nucleus where OsFD7 is predominant. Moreover, OsGF14b interacts preferably with phosphorylated OsFD7, which might be essential for the formation of a functional complex in the nuclei of the target cells. In contrast, interaction between OsFTLs and OsGF14s largely took place in the cytoplasm where OsGF14 proteins are abundantly present. Additionally, the yeast three-hybrid analysis did indeed reveal that OsFD7, OsFTLs and OsGF14s proteins interact to form a ternary complex. Together, all these results suggest that OsFD7 forms a stable complex with OsFTL and OsGF14 proteins inside the nucleus to form a functional FAC at SAM.

The domain deletion analysis revealed that the C-terminal region of OsFD7, containing GF14-3-3 binding site, is mainly responsible for its interaction with OsGF14s. An in-depth sequence analysis of motif 3 at the C-terminal end of OsFD7, predicted by SALAD database, showed the presence of a potential CDPK phosphorylation site (L-X-R/K-X-X-S) overlapping with the non-canonical GF14-3-3 binding site (R/K-X-X-S-X-S), providing clue that phosphorylation of OsFD7 could be a pre-requisite for the formation of a stable complex with OsGF14 proteins. The results of Y2H and GST pull-down experiments provided credibility to this assumption since the substitution of serine with alanine abolished interaction of OsFD7 with OsGF14b, whereas substituting serine with glutamate, a phospho-mimic substitution, restored interaction with OsGF14b.

### OsFD7 interacts directly with OsFTL proteins

Earlier reports on OsFD1 and AtFD have claimed that the interaction between FD and FT is mediated by GF14 proteins (Taoka *et al*., 2011; Kawamoto *et al*., 2015). As discussed above, we observed similar kind of result in our initial three-hybrid experiment. However, to our surprise, positive interaction was observed between OsFTLs and ΔC-ter construct of OsFD7 lacking OsGF14-3-3 binding site as well as with mutant C-terminal constructs of OsFD7 suggesting that OsFD7 has the capacity to interact with OsFTLs even in the absence of OsGF14s, which is in contrast to the previous report on OsFD1 (Taoka *et al*., 2011). The presence of non-canonical GF14-3-3 binding site in OsFD7 could possibly abrogate the requirement of OsGF14s for it to interact with OsFTLs, which is quite unique in comparison to other FDs characterized till date. One could thus speculate that the interaction of OsFD7 with OsFTLs via OsGF14-3-3 protein might be important for initiating floral transition; however, direct interaction of OsFD7 with OsFTLs could possibly explain somewhat different mode of action and additional functions it performs during rice floral initiation and development. It will be interesting to find out whether the interaction of OsFD7 with OsFTL1 under the *in vivo* conditions occurs independent of OsGF14 protein(s).

### Assembly of OsFD7-OsFTLs complex at G-box as well as C-box elements

Quite recently, Collani *et al*. (2019) and Romera-Branchat *et al*. (2020) showed that FD binds only G-box of *AP1* promoter in contrast to the previous reports claiming that bZIPs bind preferably to the ACGT DNA sequence containing conserved elements like A-box (TACGTA), C-box (GACGTC), and G-box (CACGTG) (Jakoby *et al*., 2002; Wigge *et al*., 2005; Nijhawan et al., 2008; Taoka *et al*., 2011). The binding of OsFD7 with DNA containing G-box element was confirmed by performing mobility shift assay. Additionally, a super-shifted band was observed upon the addition of OsFTL1 with OsFD7 in GB oligo gel whose migration was not affected further even after the addition of OsGF14b protein. Surprisingly, similar results were obtained using CB oligo containing C-box element. By analysing these results, one could suggest that the presence of non-canonical GF14-3-3 binding site and some additional motifs in OsFD7 might be the contributing factor for its slightly different response in comparison to other FDs identified till date in relation to its binding to both G-box as well as C-box elements. However, these results need to be further substantiated. Moreover, these results further strengthen our previous observation that OsFD7 and OsFTL1 interact directly and suggest this complex assembles at G/C-box to either activate or repress downstream target genes.

### OsCDPKs phosphorylate and activate OsFD7 in the nucleus

The presence of CDPK phosphorylation site at C-terminal end of OsFD7 and the presence of two bands in the SAM lane in the western blots gave us clue that OsFD7 could be phosphorylated at SAM by OsCDPKs. In the present study, we identified OsCDPK41 and OsCDPK49 as potential interacting partners of OsFD7 and showed that this complex is localized inside the nucleus. The nature of upper band visible in western analysis (Fig. 1C) was confirmed by an *in vivo* phosphorylation as well as de-phosphorylation by λ phosphatase treatment indicating that the extra band indeed represents the phosphorylated form of OsFD7; the phosphorylation of OsFD7 by OsCDPK49 and OsCDPK41 *in vitro* obligatorily required calcium. Although using WT and mutated (StoA substitution) C-terminal peptides of OsFD7 we could identify the motif that is phosphorylated at the serine residue *in vitro* by OsCDPK49 and OsCDPK41, but whether it is a prerequisite for the formation of FAC and onset of floral transition remains a task for future.

### OsFD7 functions in the rice flowering pathway and early stages of panicle development

Expression analysis, homology comparisons and protein-protein interaction studies involving OsFD7 provided strong evidence for its functional contribution in the rice floral transition pathway. Further, the late flowering phenotype of *OsFD7* RNAi rice transgenics substantiated that it functions as a FD or FD-like gene in rice. However, delay in the flowering time (by about 11 days) observed in these knockdown lines was not that dramatic, which could be attributed to the redundant functions the additional six FD and FD-like proteins might perform in rice flowering pathway (Taoka *et al*., 2011; Cerise *et al*., 2020). Even in an earlier study that identified OsFD1 as an interacting partner of Hd3a, the *japonica* rice RNAi lines for *OsFD1* flowered late *only* by three days (Taoka *et al*., 2011). In the present study, the down-regulation observed in the major flowering pathway genes like *OsMADS14*, *OsMADS15*, *OsMADS18* (Preston and Kellogg, 2006) and *OsMADS34* (Kobayashi *et al*., 2010; Kobayashi *et al*., 2012), in *OsFD7* RNAi lines in comparison to VC might have resulted in delayed flowering, suggesting that these MADS box genes are the likely downstream regulatory targets of OsFD7.

The *OsFD7* RNAi lines also developed denser and longer panicles with concomitant increase in the number of florets in comparison to VC plants, a unique feature not reported for other OsFDs identified from rice. In fact, the process of panicle formation is initiated at P1-1 stage, whereby branch and spikelet meristem identity is established along with the initiation of primary and secondary branches (see Nijhawan *et al*., 2008). As OsFD7 expression is high during very early stages of panicle development and declines gradually thereafter from stage P1 to P6, we could speculate that the changes observed in the panicle morphology of RNAi lines of OsFD7 could be clearly associated with the decreased expression of *OsMADS34*, which is known to control panicle development (Gao *et al*., 2010; Kobayashi *et al*., 2012). This could have resulted in the initiation of more number of primary and secondary branches as well as increase in the number of spikelets per panicle in *OsFD7* RNAi transgenics, as was also observed by Kobayashi *et al*. (2010) in the *pap2/OsMADS34* mutant of rice. Also, there was slight increase in the seed dimensions as well as in the 100 seed weight in OsFD7 RNAi plants. Therefore, it would be prudent to assume that OsFD7 fine tunes panicle architecture by acting as a negative regulator along with OsMADS34.

Although, OsFD7 downregulation gave dense inflorescence architecture but these lines displayed an increase in seed sterility with respect to VC plants. As the axis of floral meristem and organ identity is established very early by *SEP* family genes (Cui *et al*., 2010; Gao *et al*., 2010), the decrease in transcript levels of *OsMADS1*, *OsMADS7*, *OsMADS8* and *OsMADS34*, excluding *OsMADS5*, in *OsFD7* RNAi lines could have affected the development of floral organs resulting in sterility; this also signifies that OsFD7 positively regulates *SEP* family genes. However, detailed structural analysis of inflorescence and floral meristems as well as floral organs of these knockdown lines vis-à-vis VC plants would be useful in giving better insight into the precise role OsFD7 plays particularly in floral organ development.

Based on all these results, a detailed model could be proposed describing the probable mode of action of OsFD7 in flowering and panicle development (Fig. 8). OsFD7, expressing at SAM, translocates to the nucleus where its C-terminal end is phosphorylated by OsCDPK41 and OsCDPK49 resulting in the activation of OsFD7 protein. At the same time, OsFTL protein synthesized in the leaves translocates to SAM forming a complex with OsGF14 proteins in the cytoplasm. This OsFTL-OsGF14 complex enters nucleus and interacts with phosphorylated OsFD7 to form a tripartite complex, FAC, that may bind to target DNA sequences of different floral meristem identity genes triggering transition from vegetative to inflorescence meristem in rice. Surprisingly, OsFD7 interacts directly with OsFTLs in the nucleus implying that this one to one interaction is indeed providing additional stability or performing different functions during reproductive development. Apart from transition to flowering, present study shows that OsFD7 participates in regulating branch, spikelet and floral meristem development as well as floral organ development, since its knock-down lines result in more number of primary, secondary branches and unfilled seeds as well as increase in the panicle length and seed number, by directly targeting *SEP* family genes; its expression is also high during early stages of panicle development, Thus, our model depicts that OsFD7 is most likely functioning as a novel FD protein performing dual function controlling both floral transition and inflorescence development in rice, and whose function has diverged from its *Arabidopsis* counterpart AtFD as well as other rice FDs.

**Figure 8.**
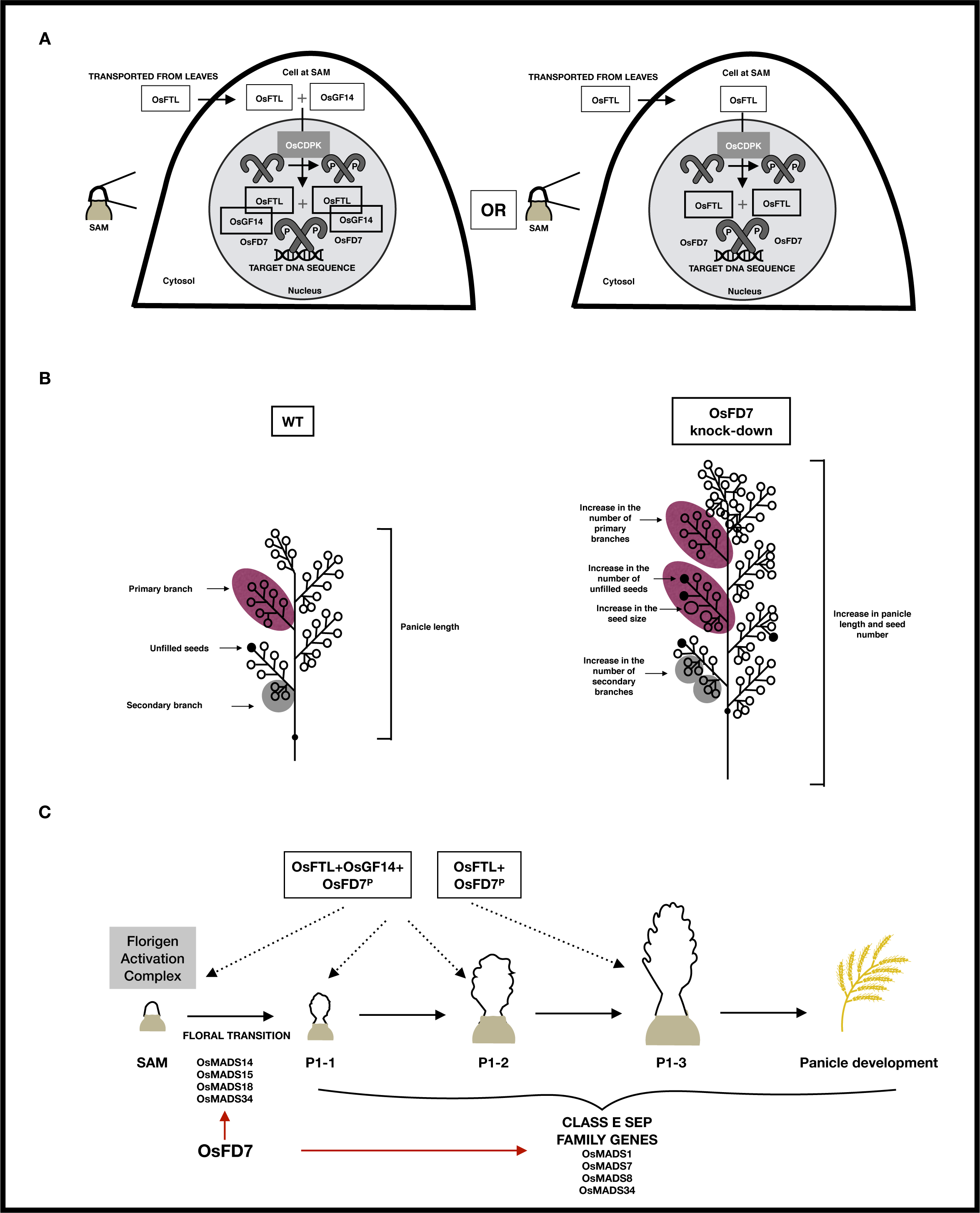
Schematic model for the involvement of OsFD7 in the formation of florigen activation complex (FAC) at SAM as well as panicle development in rice. For details, see text. Red arrow denotes positive regulation. OsFD7^P^ denotes phosphorylated form of OsFD7.

## SUPPORTING FIGURE LEGENDS

Figure S1. Genomic and protein structure of OsFD7.

Figure S2. Transactivation and localization of OsFD7.

Figure S3. Dimerization potential of OsFD7.

Figure S4. *In silico* analysis of OsFD7.

Figure S5. Expression and transactivation analysis of OsFTLs.

Figure S6. Expression analysis of OsGF14s.

Figure S7. Interaction of OsFD7 with OsFTLs and OsGF14s.

Figure S8. Interaction of OsFD7 with OsFTLs.

Figure S9. Expression and transactivation analysis of OsCDPKs.

Figure S10. Interaction of OsCDPKs and OsGF14s with truncated protein and mutated peptides of OsFD7.

Figure S11. Real-time PCR analysis of A: *OsFDs and* B: *OsCDPK41/49 and OsGF14b* genes in *OsFD7* RNAi transgenic lines with respect to VC at SAM.

Figure S12. Real-time PCR analysis of A: *OsFD1,* B: *OsFD2*, C: *OsFD3*, D: *OsFD4*, E: *OsFD6* genes in different developmental stages of rice.

Table S1. List of primers used in this study.

Table S2. List of real-time primers used in this study.

Table S3. List of plant species and their respective gene names used in this study.

Appendix S1. Supplementary text

## ACKNOWLEDGMENTS

A.K., A.N. and M.Y. thank the Council for Scientific and Industrial Research (CSIR), New Delhi, for the award of Research Fellowships. We would like to express our gratitude to Dr. A.K. Singh, Indian Agricultural Research Institute, New Delhi, for providing seed and tissue material during the course of this study.

## AUTHOR CONTRIBUTIONS

A.N. did the initial characterization of the gene; M.Y. raised rice transgenics; A.K. designed and performed the molecular interaction experiments, characterized the transgenics and prepared the manuscript with inputs from A.N. and M.Y.; J.P.K. conceived the project, supervised the work and finalized the manuscript.

## ACCESSION NUMBERS

The locus identifiers of all the genes analysed in the present study are as follows: *AtFD*/*FD-1*-AT4G35900 (AtbZIP14), *OsbZIP62*/*OsFD7*-LOC_Os07g48660, *OsFD1*-LOC_Os09g36910 (OsbZIP77), *OsFD2*-LOC_Os06g50600 (OsbZIP55), *OsFD3*-LOC_Os02g58670 (OsbZIP24), *OsFD4*-LOC_Os08g43600 (OsbZIP69), *OsFD5*-LOC_Os06g50830 (OsbZIP56), *OsFD6*-LOC_Os06g50480 (OsbZIP54), *OsFTL1*-LOC_Os01g11940, *Hd3a*-LOC_Os06g06320, *RFT1*-LOC_Os06g06300, *OsRCN1*-LOC_Os11g05470, *OsGF14b*-LOC_Os04g38870, *OsGF14c*-LOC_Os08g33370, *OsGF14d*-LOC_Os11g34450, *OsCDPK41*-LOC_Os10g41490, *OsCDPK49*-LOC_Os12g39630

## DATA AVAILABILITY STATEMENT

All data supporting the findings of this study are available within the paper and within its supplementary materials published online.

---

**Table S1.**
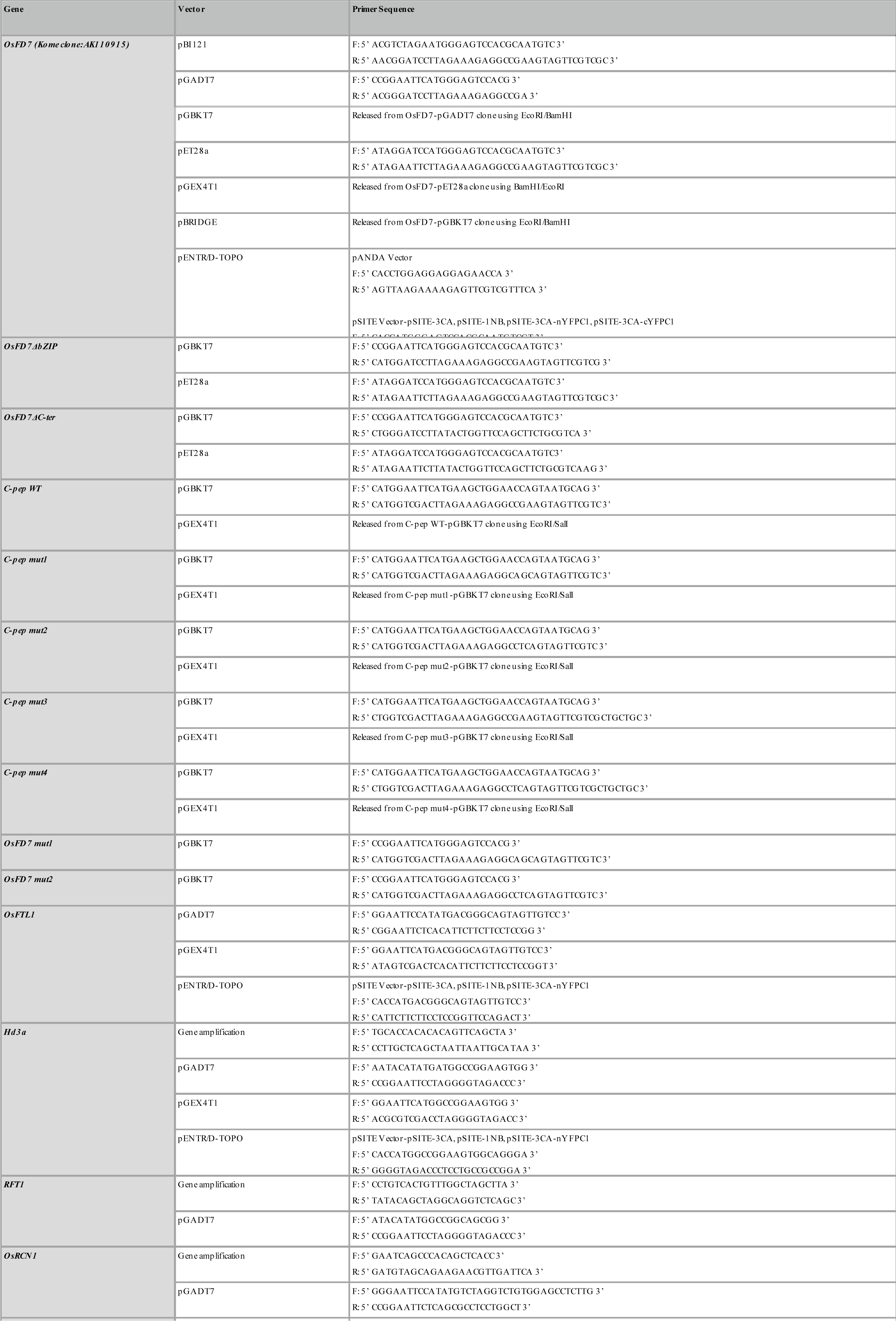
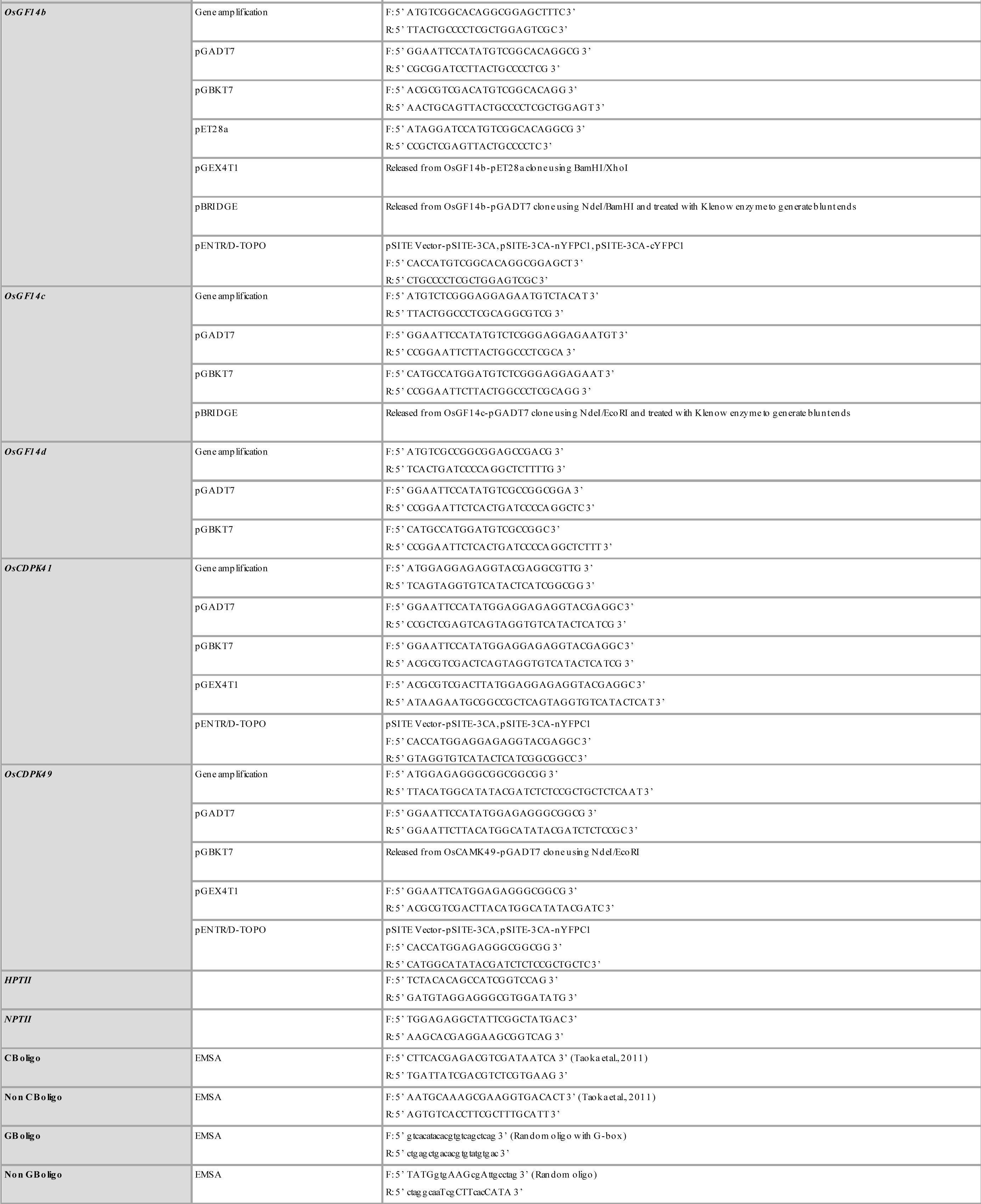
List of primers used in this study.

**Table S2.**
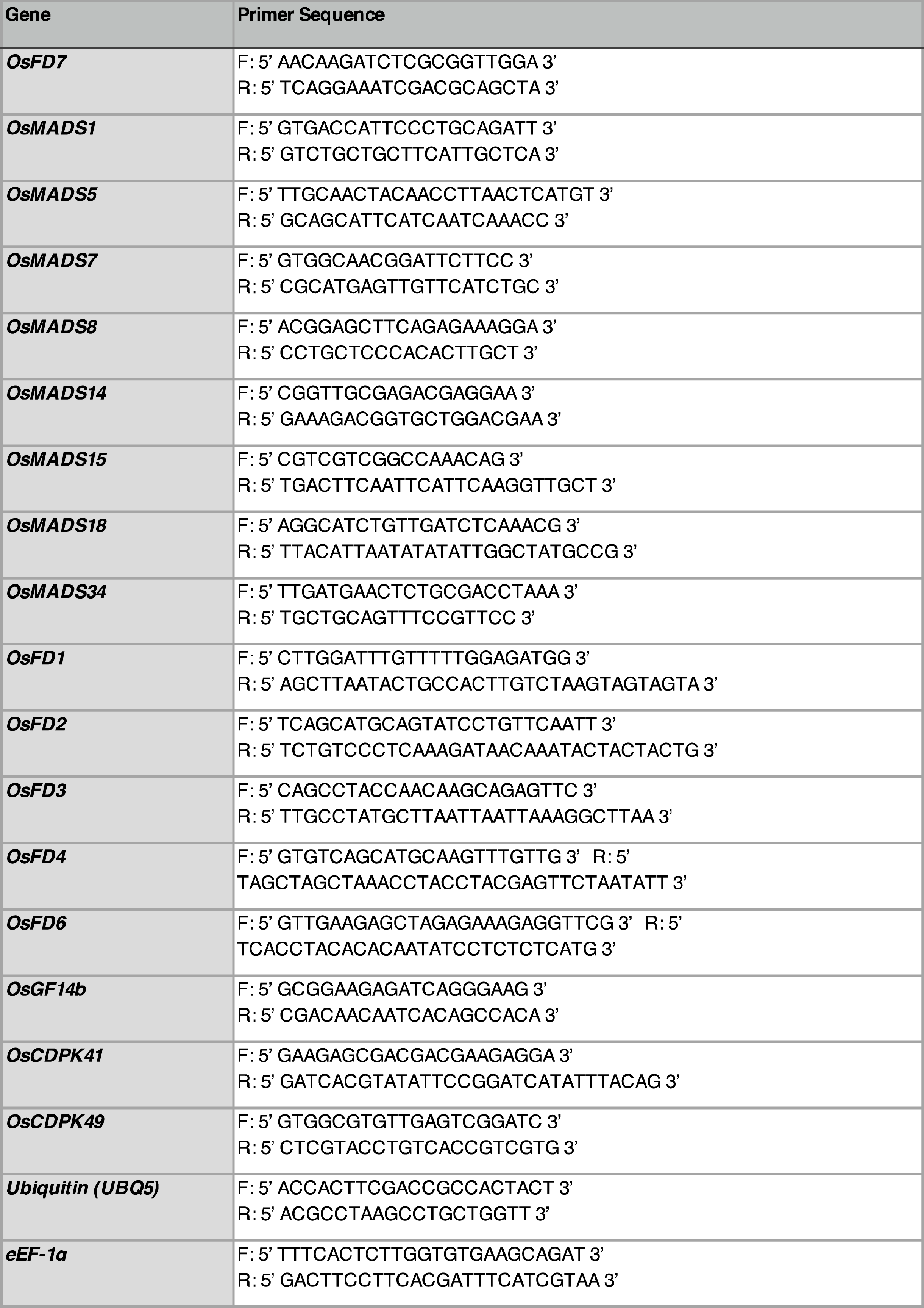
List of real-time primers used in this study.

**Table S3.**
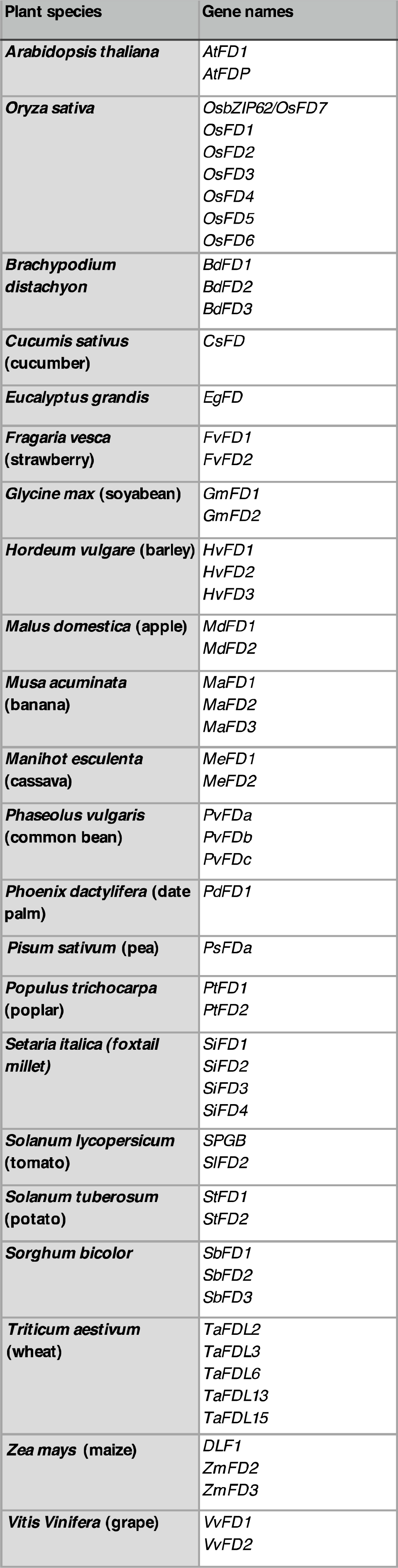
List of plant species and their respective gene names used in this study.

