## Supplemental text for "OsbZIP62/OsFD7, a functional ortholog of Flowering Locus D (FD), regulates floral transition and panicle development in rice"

**Figure S1. Genomic and protein structure of OsFD7. A:** Genomic organization of *OsFD7*. Arrows indicate start and stop codons (TIGR and KOME database). **B:** Domain organization of *OsFD7* (SMART database)(Schultz et al., 1998). **C:** 3D Robetta modelling of OsFD7 comprising of α helical structure. Basic leucine-zipper domain highlighted with blue colour and C-terminal coiled-coil region highlighted with green colour (Kim et al., 2004) (<http://www.jmol.org/>).

**Figure S2. Transactivation and localization of OsFD7. A:** Transactivation activity assay of OsFD7 protein in yeast. Positive control: OsbZIP16-pGBKT7; Vector control: pGBKT7 vector. BD indicates DNA binding domain of GAL4. -HW: selective medium (SD/-His-Trp); -W: control medium (SD/-Trp). **B:** Comparison of the putative NLS sequence of OsZIP62 with ZmO2 and OsFD1. Amino acids critical for nuclear targeting are highlighted by colored boxes. **C:** Sub-cellular localization of OsFD7 protein in onion peel cells. Empty pSITE-3CA vector used as vector control.

**Figure S3. Dimerization potential of OsFD7. A:** Region responsible for dimerization specificity and stability is highlighted with blue and pink color. The predicted C- terminal boundary is denoted by the symbol # (Nijhawan et al., 2008). **B:** Y2H assay showing homo-dimerization of OsFD7. Positive control: pGADT7-T + pGBKT7-53; Negative control: pGADT7-T + pGBKT7-Lam. AD and BD denotes activation domain and DNA binding domain of GAL4, respectively. -HLW+3-AT: selective medium (SD/-His-Leu-Trp) supplemented with 3-AT; -LW: control medium (SD/-Leu-Trp). **C:** GST pull-down assay confirming homo-dimerization of OsFD7. OsFD7-His: His tagged OsFD7 protein; OsFD7-GST: GST tagged OsFD7 protein; GST: GST protein. **D:** In BiFC assay, homo-dimerization seen in the nucleus. Negative control: pSITE nYFPC1+pSITE cYFPC1. **E:** Diagrammatic representation of full length and truncated OsFD7 proteins. **F:** Y2H assay using truncated OsFD7. Positive control: pGADT7-T + pGBKT7-53; Negative control: pGADT7-T + pGBKT7-Lam. AD and BD denotes activation domain and DNA binding domain of GAL4, respectively. -HLW+3-AT: selective medium (SD/-His-Leu-Trp) supplemented with 3-AT; -LW: control medium (SD/-Leu-Trp). **G:** GST pull-down assay using truncated OsFD7. ΔbZIP-His: truncated OsFD7 His tagged protein; ΔC-ter-His: C-terminal truncated OsFD7 His tagged protein; OsFD7-GST: GST tagged OsFD7 protein; GST: GST protein.

**Figure S4. *In silico* analysis of OsFD7. A:** Conserved amino acid motifs and their combinations predicted in the protein sequences of FD and FD-like proteins using the SALAD database (Mihara et al. 2010). **B:** MAFFT alignment of the FD and FD-like proteins highlighting basic region, leucine-zipper region, GF14-3-3 recognition/CDPK phosphorylation site and SAP motif conserved in almost all the species. **C:** Phylogenetic analysis of *OsFD7* with FD and FD-like proteins across various genera.

**Figure S5. Expression and transactivation analysis of OsFTLs. A:** Heat maps depicting expression profile of *OsFD7* and *OsFTLs* during different stages of rice development (Rice oligonucleotide array database). **B:** Transactivation activity assay of OsFTLs and OsRCN1 protein in yeast. Positive control: OsbZIP16-pGBKT7; Vector control: pGBKT7 vector. BD indicates DNA binding domain of GAL4. -HW: selective medium (SD/-His-Trp); -W: control medium (SD/-Trp). **C:** Sub-cellular localization of OsFTL1 and Hd3a protein in onion peel cells. Vector control: pSITE-3CA vector.

**Figure S6. Expression analysis of OsGF14s. A:** Heat maps depicting expression profile of OsFD7 and OsGF14s during different stages of rice development (Rice oligonucleotide array database). **B:** Sub-cellular localization of OsGF14b protein in onion peel cells. Vector control: pSITE-3CA vector.

**Figure S7. Interaction of OsFD7 with OsFTLs and OsGF14s. A:** Y2H analysis of OsFD7, OsGF14s and OsFTLs. Positive control: pGADT7-T + pGBKT7-53; Negative control: pGADT7-T + pGBKT7-Lam. AD and BD denotes activation domain and DNA binding domain of GAL4, respectively. -HLW+3-AT: selective medium (SD/-His-Leu-Trp) supplemented with 3-AT; -LW: control medium (SD/-Leu-Trp). **B:** GST pull-down assay confirming interactions among OsFD7, OsFTLs and OsGF14b. OsFD7-His: His tagged OsFD7 protein; OsGF14b-His: His tagged OsGF14b protein; OsGF14b-GST: GST tagged OsGF14b protein; OsFTL1-GST and Hd3a-GST: GST tagged OsFTL1 and Hd3a protein; GST: GST protein. **C:** BiFC assay showing interaction between OsFD7 and OsGF14b proteins in the nucleus and that of OsFTL1 with OsGF14b in the cytoplasm of onion peel cells. Negative controls: OsFTL1+OsbZIP16 and OsGF14b+OsbZIP16. **D:** Yeast three hybrid system (modified yeast two hybrid system) demonstrating the three-way interaction among OsFD7, OsGF14s and OsFTLs proteins using pBRIDGE vector (Clonetech). See Material and Methods and text in Results for details.

**Figure S8. Interaction of OsFD7 with OsFTLs. A:** GST pull-down assay between OsFTLs and purified OsFD7 proteins. OsFD7 purified His: OsFD7 His tagged protein purified using Ni-NTA column; OsFTL1-GST: GST tagged OsFLT1 protein; Hd3a-GST: GST tagged Hd3a protein; GST: GST protein. **B:** DNA mobility shift assay to determine interaction of C-box element with different components. OsFD7GST: GST tagged OsFD7 protein; OsGF14bGST: GST tagged OsGF14b protein; OsFTL1GST: GST tagged OsFTL1 protein; ALL: OsFD7GST + OsFTL1GST+ OsGF14bGST; GST: GST protein. CB oligo: C-box element oligo; Non CB oligo: Non C-box element oligo. DNA mobility shifts marked with red arrows.

**Figure S9. Expression and transactivation analysis of OsCDPKs. A:** Putative **i**nteracting partners of OsFD7 predicted by STRING database (Szklarczyk et al., 2015). **B:** Heat maps depicting expression profile of OsFD7 and OsCDPKs during different stages of rice development (Rice oligonucleotide array database). **C:** Transactivation property of OsCDPKs tested using Y2H assay. Positive control: OsbZIP16-pGBKT7; Vector control: pGBKT7 vector. BD indicates DNA binding domain of GAL4. -HW+3-AT: selective medium (SD/-His-Trp) supplemented with 3-AT; -W: control medium (SD/-Trp). **D:** Sub-cellular localization of OsCDPKs in onion peel cells. Vector control: pSITE-3CA vector.

**Figure S10. Interaction of OsCDPKs and OsGF14s with truncated protein and mutated peptides of OsFD7. A:** Y2H assay depicting protein-protein interactions using truncated OsFD7. Positive control: pGADT7-T + pGBKT7-53; Negative control: pGADT7-T + pGBKT7-Lam. AD and BD denotes activation domain and DNA binding domain of GAL4, respectively. -HLW+3-AT: selective medium (SD/-His-Leu-Trp) supplemented with 3-AT; -LW: control medium (SD/-Leu-Trp). **B:** GST pull-down assay of OsCDPKs with truncated OsFD7. ΔbZIP-His: truncated OsFD7 His tagged protein; ΔC-ter-His: truncated OsFD7 His tagged protein; OsCDPK41,49-GST: GST tagged OsCDPK41,49 protein; GST: GST protein. **C:** Diagrammatic representation of mutated peptide of OsFD7 harbouring amino acid substitutions. **D:** Y2H assay using truncated OsFD7 and mutated OsFD7 peptides. Positive control: pGADT7-T + pGBKT7-53; Negative control: pGADT7-T + pGBKT7-Lam. AD and BD denotes activation domain and DNA binding domain of GAL4, respectively. -HLW+3-AT: selective medium (SD/-His-Leu-Trp) supplemented with 3-AT; -LW: control medium (SD/-Leu-Trp). **E:** GST pull-down assay performed with mutated peptides of OsFD7. C-pep mut3,mut4-GST: GST tagged mutated OsFD7 C terminal peptides; GST: GST protein. **F:** *In vitro* kinase assay using GST tagged C-terminal peptide of OsFD7 by OsCDPK49 using anti-phosphoserine. No band could be seen with mutated C-terminal peptide of OsFD7. OsFD7 C-pep mut3-GST: OsFD7 C-terminal GST tagged peptide with a single amino acid substitution LtoQ; OsCDPK49-GST: GST tagged OsCDPK49 protein; GST: GST protein.

**Figure S11. Real-time PCR analysis of** **A:** *OsFDs and* **B:** *OsCDPK41/49 and OsGF14b* genes in *OsFD7* RNAi transgenic lines with respect to VC at SAM. *UBQ5* and *eEF-1α* genes were used as internal controls (Jain et al., 2006). Error bars represent standard error.

**Figure S12. Real-time PCR analysis of** **A:** *OsFD1,* **B:** *OsFD2*, **C:** *OsFD3*, **D:** *OsFD4*, **E:** *OsFD6 genes* in different developmental stages of rice. *UBQ5* and *eEF-1α* genes were used as internal controls (Jain et al., 2006). Error bars represent standard error.

**Table S1. List of primers used in this study.**

**Table S2. List of real-time primers used in this study.**

**Table S3. List of plant species and their respective gene names used in this study.**

**Appendix S1. Supplementary text.**

**Appendix S1-** **Supplementary text**

**OsFD7 harbours a conserved bZIP domain and forms an α-helical structure**

*OsFD7* gene encodes a protein of 274 aa residues containing a well conserved bZIP domain (179 to 248 aa) towards its C-terminus as predicted by SMART domain analysis software. This bZIP domain comprises of an invariant motif (N-X_7_-R/K) in the basic region and heptad repeats in the leucine-zipper region, which is approximately 9 aa downstream of R/K residue, essential for the sequence specific DNA-binding as well as dimerization activity (Fig. S1B). For examining the transactivation potential of OsFD7, its CDS was cloned in pGBKT7 vector. OsFD7 does not cause transactivation on its own as no colonies appeared on the selection medium in Y2H analysis (Fig. S2A); OsbZIP16 was used as a positive control (Chen *et al*., 2012; Pandey *et al*., 2018), Although, a number of bZIPs harbor an additional activation domain required to activate transcription of the downstream targets (Schindler *et al*., 1992; Chen *et al*., 2012), however, OsFD7 lacks this transactivation capacity and, therefore, could possibly work as part of a complex to regulate transcription of its target genes. In addition, 3D structure prediction by Robetta server showed that OsFD7 is organized into a α-helical structure (Fig. S1C) having significant similarity with yeast GCN4 protein, which is stabilized by its C-terminal coiled-coil region. This C-terminal coiled-coil region of bZIPs stabilizes the dimerization interface for the formation of a successful homo- or hetero-dimers (Ellenberger *et al*., 1992).

**OsFD7 protein is nuclear localized and forms a stable homodimer in the nucleus**

PSORT prediction software identified a putative bipartite nuclear localization signal (NLS) within the basic region of bZIP domain (179 to 209 aa). This NLS sequence of OsFD7 is very similar to OsFD1 of *Oryza sativa* (Tsuji *et al*., 2013) and Opaque-2 of *Zea mays* (Varagona and Raikhel, 1994) (Fig. S2B). To determine the subcellular localization of OsFD7, its CDS was cloned in pSITE 3CA vector and particle bombardment of fresh onion peel cells was carried out. YFP fluorescence (representing OsFD7 protein) was detected exclusively in the nucleus (Fig. S2C). By analysing these results, we could state that OsFD7 is indeed nuclear localized, providing evidence that NLS is functional.

Further, Nijhawan *et al*. (2008) classified bZIPs into several subfamilies. OsFD7 belonged to BZ1 subfamily containing a dimerizing domain (Fig. S3A). Since the bZIP proteins have the capacity to form homo-dimers, which is essential to make contact with the DNA molecule at an appropriate position and regulate transcription (Landschulz *et al*., 1988), the dimerizing nature of OsFD7 could be confirmed experimentally by yeast-2-hybrid (Y2H) and GST pull-down assays (Fig. S3B; Fig. S3C). In bimolecular fluorescence complementation (BiFC) assay, YFP fluorescence was detected preferentially in the nucleus of onion peel cells (Fig. S3D) where OsFD7 primarily localizes, thus providing sufficient evidence for its homo-dimerizing capability. In order to substantiate the involvement of bZIP domain in homo-dimerization of OsFD7, two deletion constructs were generated: ΔbZIP (bZIP domain deletion construct: 179 to 248 aa) and ΔC-ter (C-terminal region deletion construct: 249 to 274 aa) (Fig. S3E). Both Y2H and GST pull-down assays showed no interaction when ΔbZIP construct was used whereas ΔC-ter construct gave positive interaction (Fig. S3F; Fig. S3G) suggesting that even though bZIP domain is mainly required for homo-dimerization but C-terminal coiled-coil region may be required for a more stable homo-dimer; however, further studies are required to substantiate this assumption.

**Direct interaction of OsFD7 with OsFTLs**

To rule out the possibility of any bacterial accessory factors mediating OsFD7-OsFTLs interaction, we performed GST pull-down assay using purified OsFD7 protein. In concurrence with the previous results, a direct interaction was observed in this case as well (Fig. S8A).

**OsFD7 interacts with calcium dependent protein kinases (OsCDPKs)**

To find out which region of OsFD7 is crucial for its interaction with OsCDPKs, ΔbZIP, ΔC-ter and C-pep WT constructs of OsFD7 were used for performing Y2H and GST pull-down assays. All these OsFD7 deletion constructs slightly affected its interaction with OsCDPKs in Y2H experiment suggesting that OsCDPK41,49 interact with OsFD7 not only at its C-terminal end but at other positions as well (Fig. S10A, B).

As CDPKs require the presence of leucine residue, a hydrophobic amino acid, at -5 position for it to recognize and phosphorylate serine/threonine residue within L-X-R/K-X-X-S/T sequence (Vlad *et al*., 2008), therefore, mutation of this residue would abolish phosphorylation of OsFD7 by OsCDPKs and might affect its interaction with OsGF14s since OsGF14s recognize and interact with phosphorylated OsFD7 (Fig. S7B). To validate this assumption, two mutant constructs were generated: one in which leucine was substituted with glutamine (named as C-pep m3-LtoQ) while the other mutant construct contains a serine to glutamate substitution in the LtoQ background (named as C-pep m4-StoE in LtoQ) (Fig. S10C). In Y2H analysis, C-pep m3 displayed weak interaction with OsGF14b while this interaction was fully restored with C-pep m4 mutant harbouring phospho-mimic substitution (Fig. S10D). Similarly, GST pull down assay performed using these mutant peptides showed no interaction with C-pep m3 whereas strong signal was observed in the case of C-pep m4 mutant (Fig. S10E). These results suggest that L-R-R-T-T-S sequence at the C-terminal end of OsFD7 strictly follows this rule implying that OsCDPKs phosphorylates S271 residue of OsFD7. To further confirm the requirement of leucine for OsCDPK activity, an *in vitro* kinase assay was carried out in the presence of calcium with the purified C-pep m3-LtoQ peptide using OsCDPK49 purified protein. No band could be detected after western blotting in the 5^th^ lane containing purified C-pep m3-LtoQ (LtoQ substitution) peptide of OsFD7 (Fig. S10F) confirms that leucine at -5 position is critical for the phosphorylation of S271 of OsFD7 and subsequently for its interaction with OsGF14s.

**Specificity of RNAi construct**

The previous reports by Taoka *et al*. (2011) and Tsuji *et al*. (2013) claimed the presence of six *FD* genes in rice. In order to demonstrate the specificity of this RNAi construct, expression levels of earlier identified *FDs* were also checked in these lines. Their transcript levels were similar in both VC and transgenics (Fig. S11), indicating that delayed flowering phenotype is not an off-target effect. Also, protein-protein interaction assays carried out earlier showed a direct interaction of OsFD7 with OsGF14s as well as OsCDPKs selected for this study. The mRNA levels of *OsGF14b*, *OsCDPK41* and *OsCDPK49* also remained unaffected in these RNAi transgenics (Fig. S11B).
