## Supplemental figures for "OsbZIP62/OsFD7, a functional ortholog of Flowering Locus D (FD), regulates floral transition and panicle development in rice"

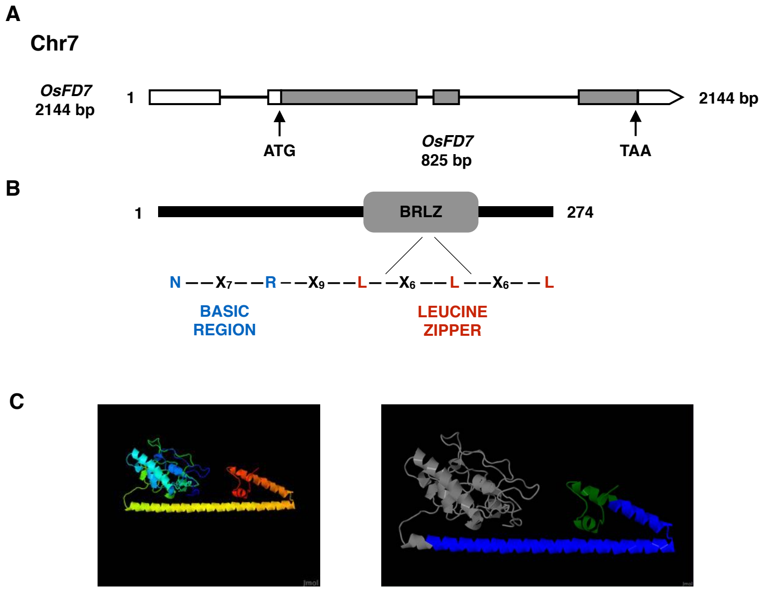


**Figure S1. Genomic and protein structure of OsFD7. A:** Genomic organization of *OsFD7*. Arrows indicate start and stop codons (TIGR and KOME database). **B:** Domain organization of *OsFD7* (SMART database)(Schultz et al., 1998). **C:** 3D Robetta modelling of OsFD7 comprising of α helical structure. Basic leucine-zipper domain highlighted with blue colour and C-terminal coiled-coil region highlighted with green colour (Kim et al., 2004) (http://www.jmol.org/).

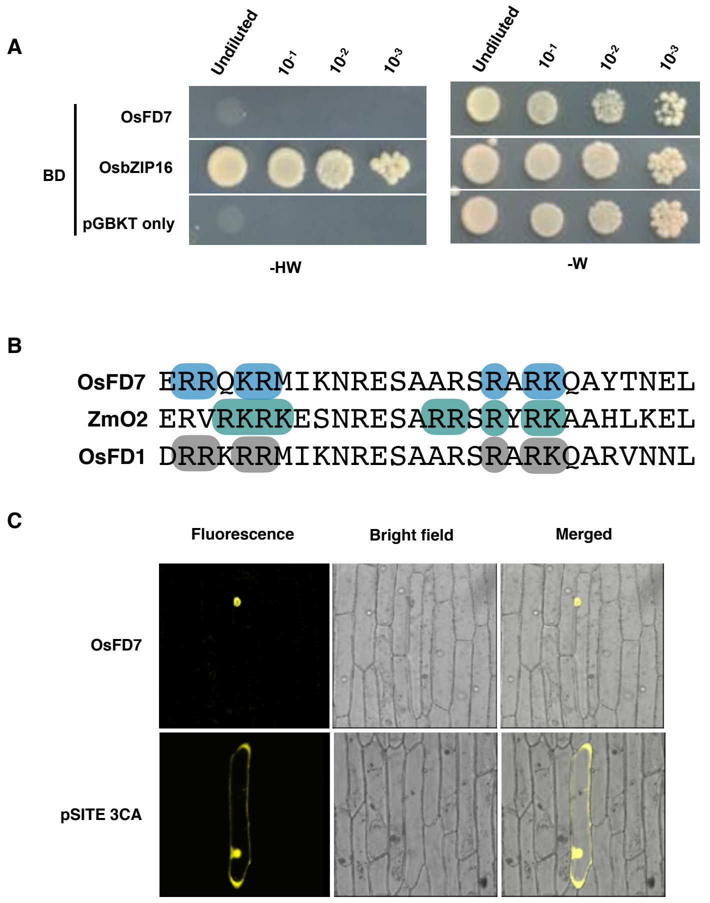


**Figure S2. Transactivation and localization of OsFD7. A:** Transactivation activity assay of OsFD7 protein in yeast. Positive control: OsbZIP16-pGBKT7; Vector control: pGBKT7 vector. BD indicates DNA binding domain of GAL4. -HW: selective medium (SD/-His-Trp); -W: control medium (SD/-Trp). **B:** Comparison of the putative NLS sequence of OsZIP62 with ZmO2 and OsFD1. Amino acids critical for nuclear targeting are highlighted by colored boxes. **C:** Sub-cellular localization of OsFD7 protein in onion peel cells. Empty pSITE-3CA vector used as vector control.

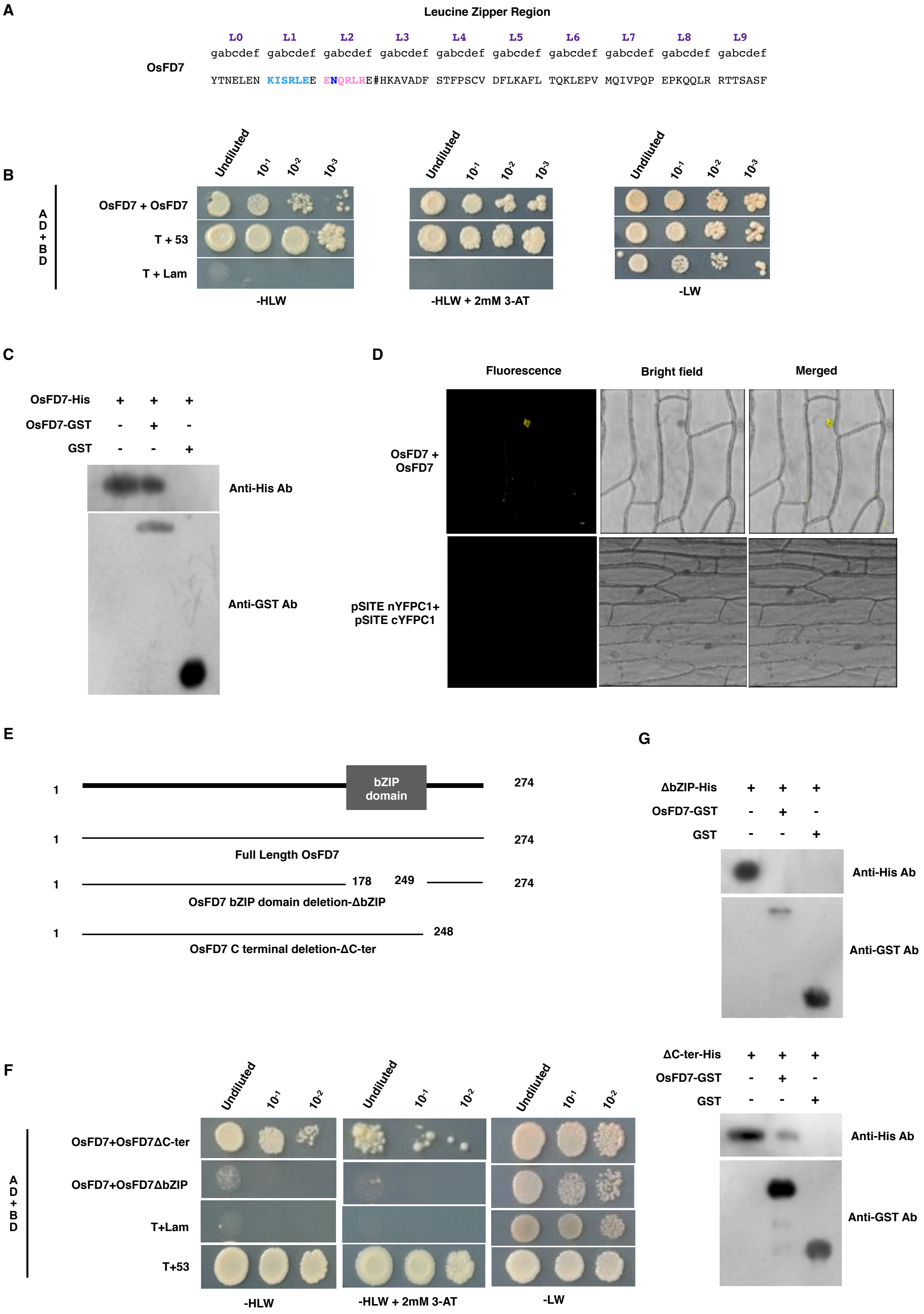


**Figure S3. Dimerization potential of OsFD7. A:** Region responsible for dimerization specificity and stability is highlighted with blue and pink color. The predicted C- terminal boundary is denoted by the symbol # (Nijhawan et al., 2008). **B:** Y2H assay showing homo-dimerization of OsFD7. Positive control: pGADT7-T + pGBKT7-53; Negative control: pGADT7-T + pGBKT7-Lam. AD and BD denotes activation domain and DNA binding domain of GAL4, respectively. -HLW+3-AT: selective medium (SD/-His-Leu-Trp) supplemented with 3-AT; -LW: control medium (SD/-Leu-Trp). **C:** GST pull-down assay confirming homo-dimerization of OsFD7. OsFD7-His: His tagged OsFD7 protein; OsFD7-GST: GST tagged OsFD7 protein; GST: GST protein. **D:** In BiFC assay, homo-dimerization seen in the nucleus. Negative control: pSITE nYFPC1+pSITE cYFPC1. **E:** Diagrammatic representation of full length and truncated OsFD7 proteins. **F:** Y2H assay using truncated OsFD7. Positive control: pGADT7-T + pGBKT7-53; Negative control: pGADT7-T + pGBKT7-Lam. AD and BD denotes activation domain and DNA binding domain of GAL4, respectively. -HLW+3-AT: selective medium (SD/-His-Leu-Trp) supplemented with 3-AT; -LW: control medium (SD/-Leu-Trp). **G:** GST pull-down assay using truncated OsFD7. ΔbZIP-His: truncated OsFD7 His tagged protein; ΔC-ter-His: C-terminal truncated OsFD7 His tagged protein; OsFD7-GST: GST tagged OsFD7 protein; GST: GST protein.

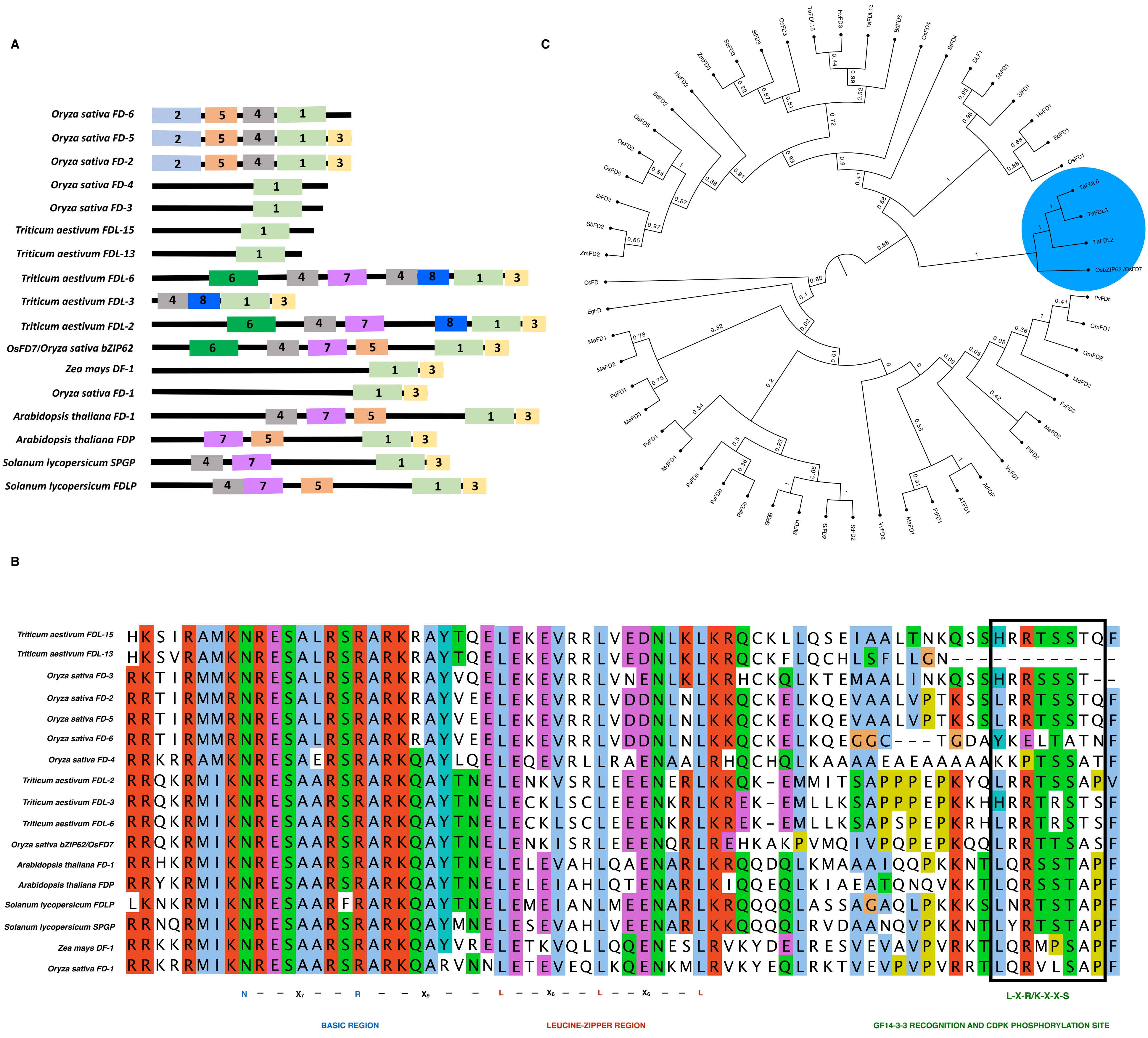


**Figure S4.** ***In silico*** **analysis of OsFD7. A:** Conserved amino acid motifs and their combinations predicted in the protein sequences of FD and FD-like proteins using the SALAD database (Mihara et al. 2010). **B:** MAFFT alignment of the FD and FD-like proteins highlighting basic region, leucine-zipper region, GF14-3-3 recognition/CDPK phosphorylation site and SAP motif conserved in almost all the species. **C:** Phylogenetic analysis of *OsFD7* with FD and FD-like proteins across various genera.

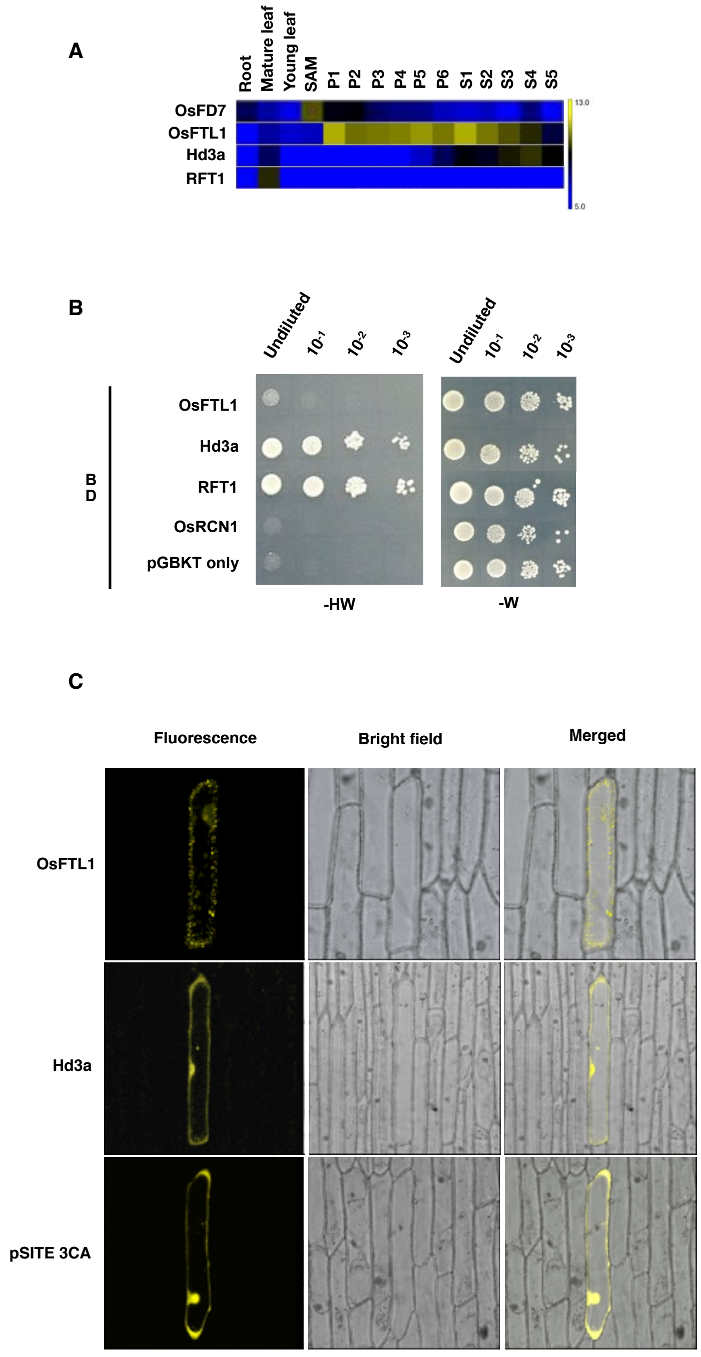


**Figure S5. Expression and transactivation analysis of OsFTLs. A:** Heat maps depicting expression profile of *OsFD7* and *OsFTLs* during different stages of rice development (Rice oligonucleotide array database). **B:** Transactivation activity assay of OsFTLs and OsRCN1 protein in yeast. Positive control: OsbZIP16-pGBKT7; Vector control: pGBKT7 vector. BD indicates DNA binding domain of GAL4. -HW: selective medium (SD/-His-Trp); -W: control medium (SD/-Trp). **C:** Sub-cellular localization of OsFTL1 and Hd3a protein in onion peel cells. Vector control: pSITE-3CA vector.

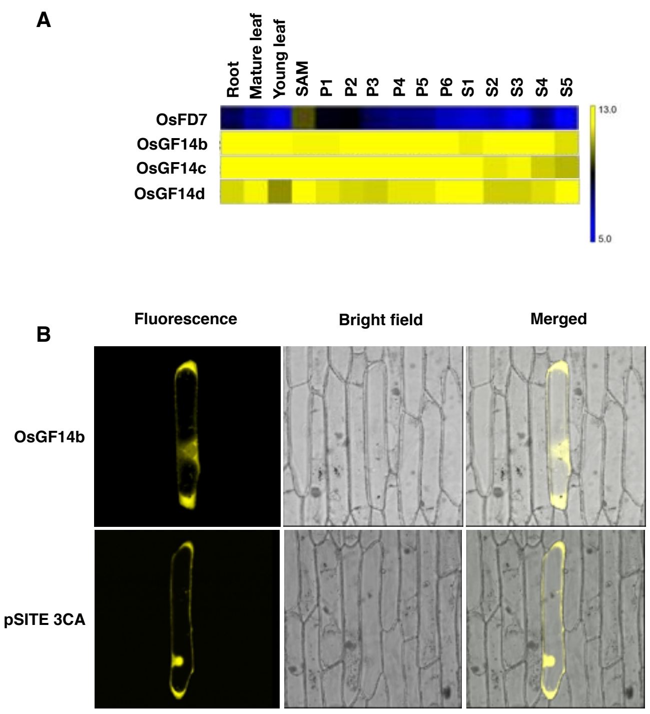


**Figure S6. Expression analysis of OsGF14s. A:** Heat maps depicting expression profile of OsFD7 and OsGF14s during different stages of rice development (Rice oligonucleotide array database). **B:** Sub-cellular localization of OsGF14b protein in onion peel cells. Vector control: pSITE-3CA vector.

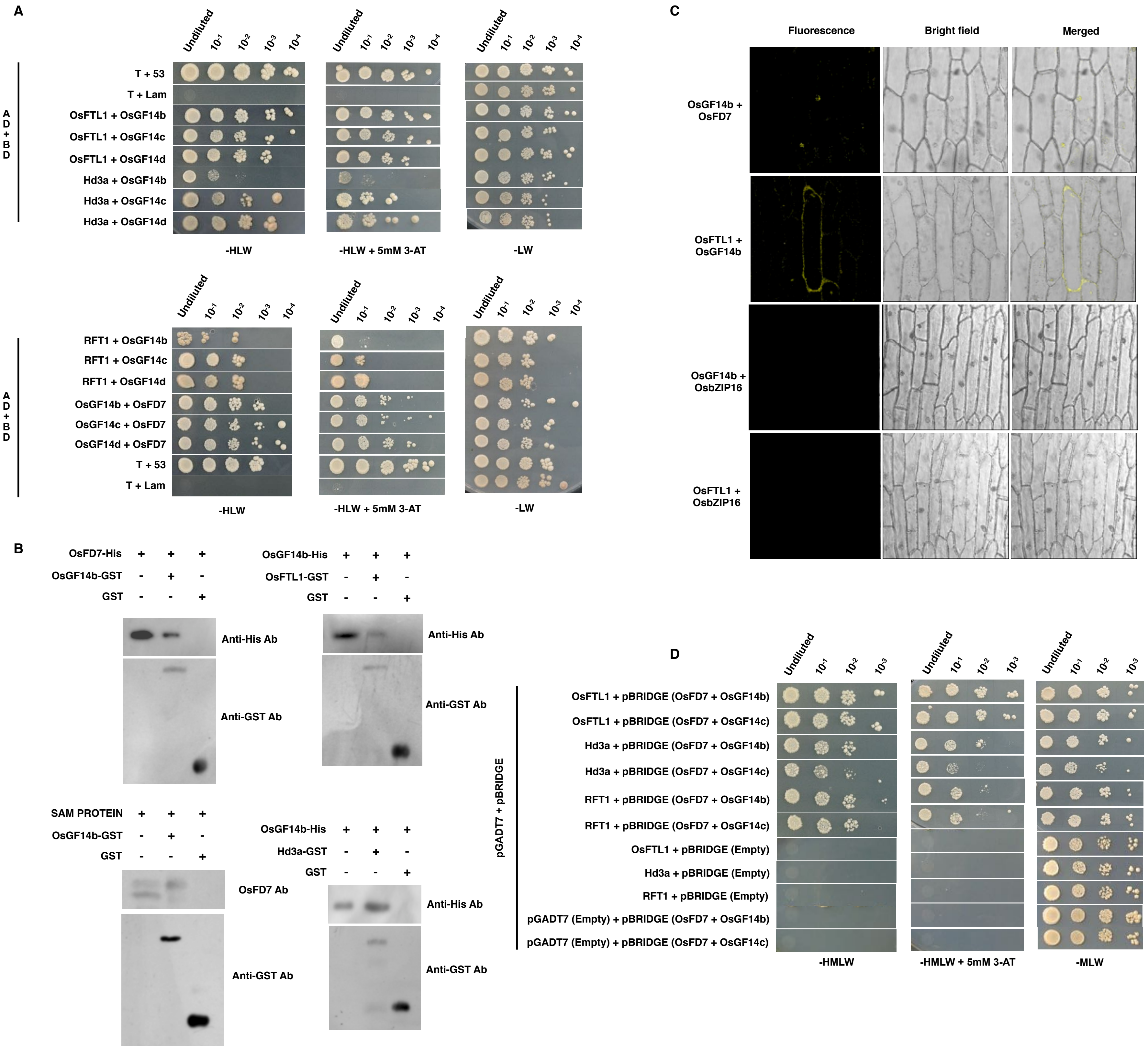


**Figure S7. Interaction of OsFD7 with OsFTLs and OsGF14s. A:** Y2H analysis of OsFD7, OsGF14s and OsFTLs. Positive control: pGADT7-T + pGBKT7-53; Negative control: pGADT7-T + pGBKT7-Lam. AD and BD denotes activation domain and DNA binding domain of GAL4, respectively. -HLW+3-AT: selective medium (SD/-His-Leu-Trp) supplemented with 3-AT; -LW: control medium (SD/-Leu-Trp). **B:** GST pull-down assay confirming interactions among OsFD7, OsFTLs and OsGF14b. OsFD7-His: His tagged OsFD7 protein; OsGF14b-His: His tagged OsGF14b protein; OsGF14b-GST: GST tagged OsGF14b protein; OsFTL1-GST and Hd3a-GST: GST tagged OsFTL1 and Hd3a protein; GST: GST protein. **C:** BiFC assay showing interaction between OsFD7 and OsGF14b proteins in the nucleus and that of OsFTL1 with OsGF14b in the cytoplasm of onion peel cells. Negative controls: OsFTL1+OsbZIP16 and OsGF14b+OsbZIP16. **D:** Yeast three hybrid system (modified yeast two hybrid system) demonstrating the three-way interaction among OsFD7, OsGF14s and OsFTLs proteins using pBRIDGE vector (Clonetech). See Material and Methods and text in Results for details.

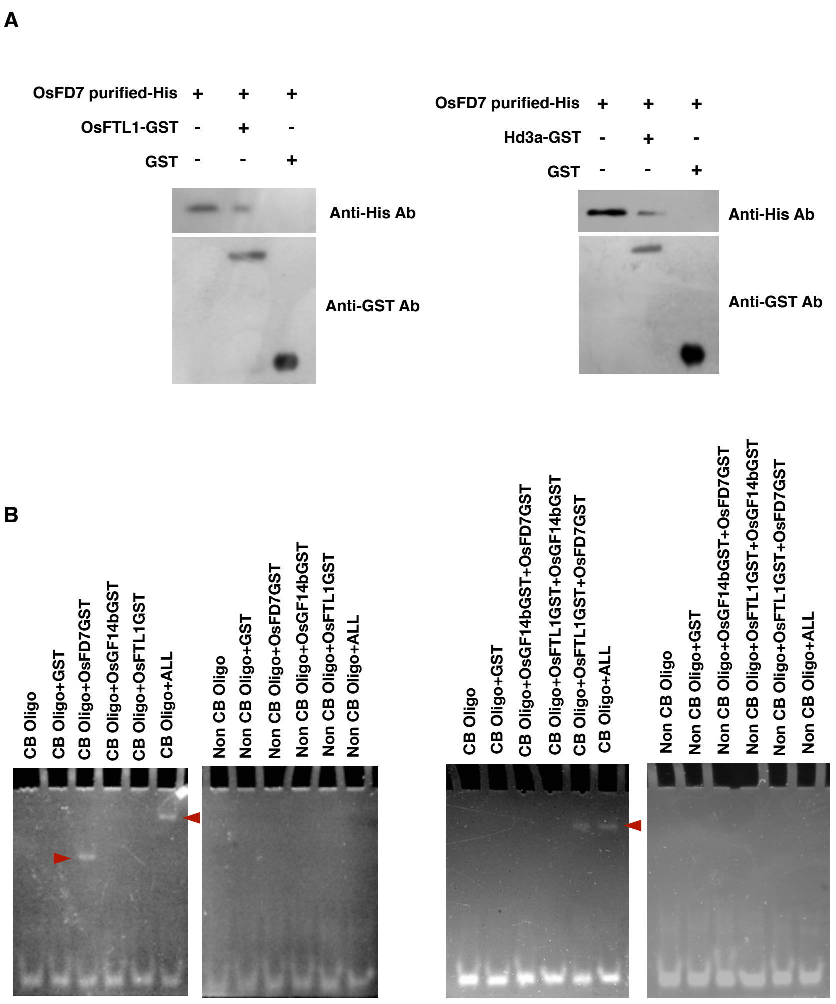


**Figure S8. Interaction of OsFD7 with OsFTLs. A:** GST pull-down assay between OsFTLs and purified OsFD7 proteins. OsFD7 purified His: OsFD7 His tagged protein purified using Ni-NTA column; OsFTL1-GST: GST tagged OsFLT1 protein; Hd3a-GST: GST tagged Hd3a protein; GST: GST protein. **B:** DNA mobility shift assay to determine interaction of C-box element with different components. OsFD7GST: GST tagged OsFD7 protein; OsGF14bGST: GST tagged OsGF14b protein; OsFTL1GST: GST tagged OsFTL1 protein; ALL: OsFD7GST + OsFTL1GST+ OsGF14bGST; GST: GST protein. CB oligo: C-box element oligo; Non CB oligo: Non C-box element oligo. DNA mobility shifts marked with red arrows.

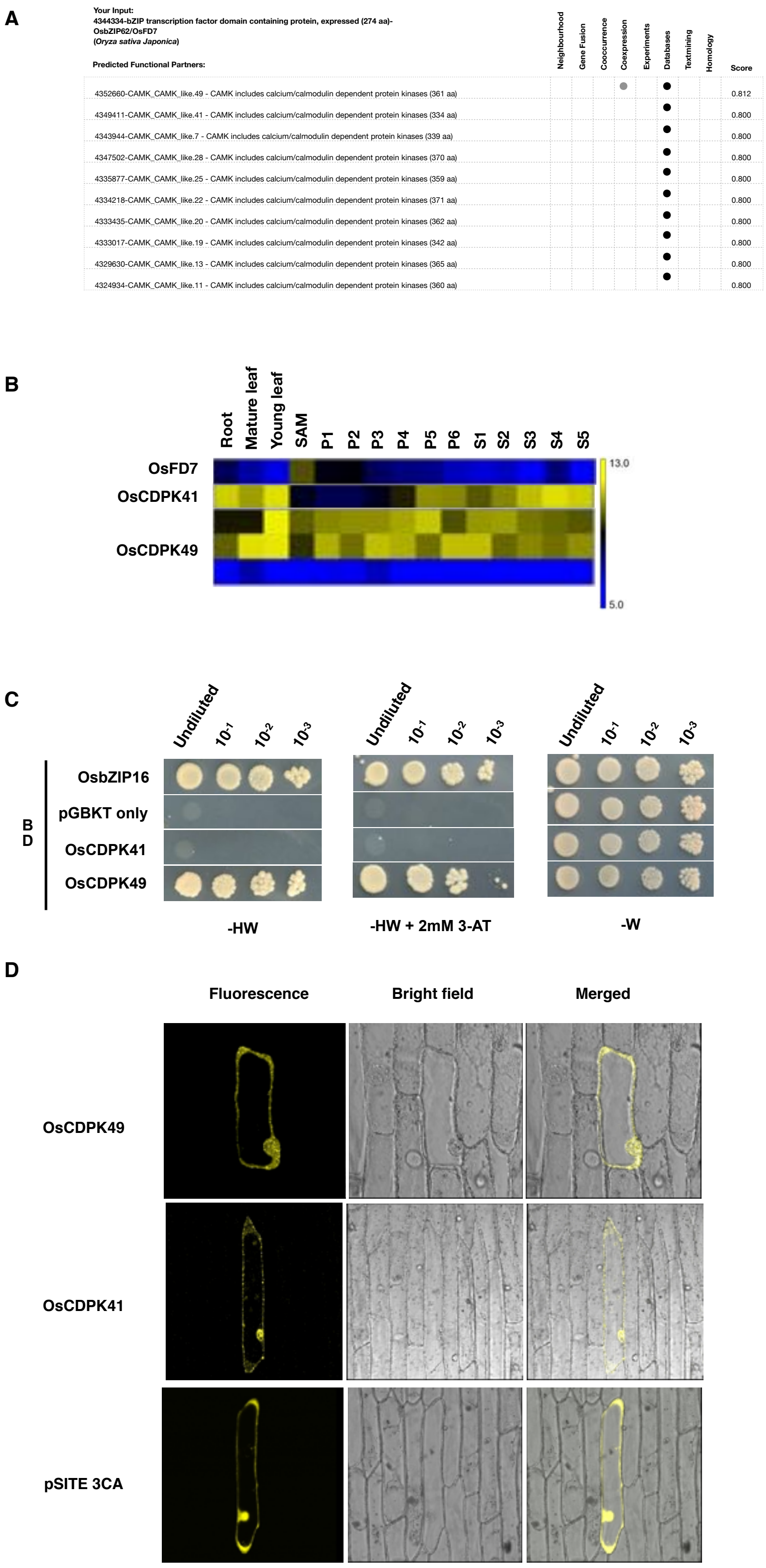


**Figure S9. Expression and transactivation analysis of OsCDPKs. A:** Putative **i**nteracting partners of OsFD7 predicted by STRING database (Szklarczyk et al., 2015). **B:** Heat maps depicting expression profile of OsFD7 and OsCDPKs during different stages of rice development (Rice oligonucleotide array database). **C:** Transactivation property of OsCDPKs tested using Y2H assay. Positive control: OsbZIP16-pGBKT7; Vector control: pGBKT7 vector. BD indicates DNA binding domain of GAL4. -HW+3-AT: selective medium (SD/-His-Trp) supplemented with 3-AT; -W: control medium (SD/-Trp). **D:** Sub-cellular localization of OsCDPKs in onion peel cells. Vector control: pSITE-3CA vector.

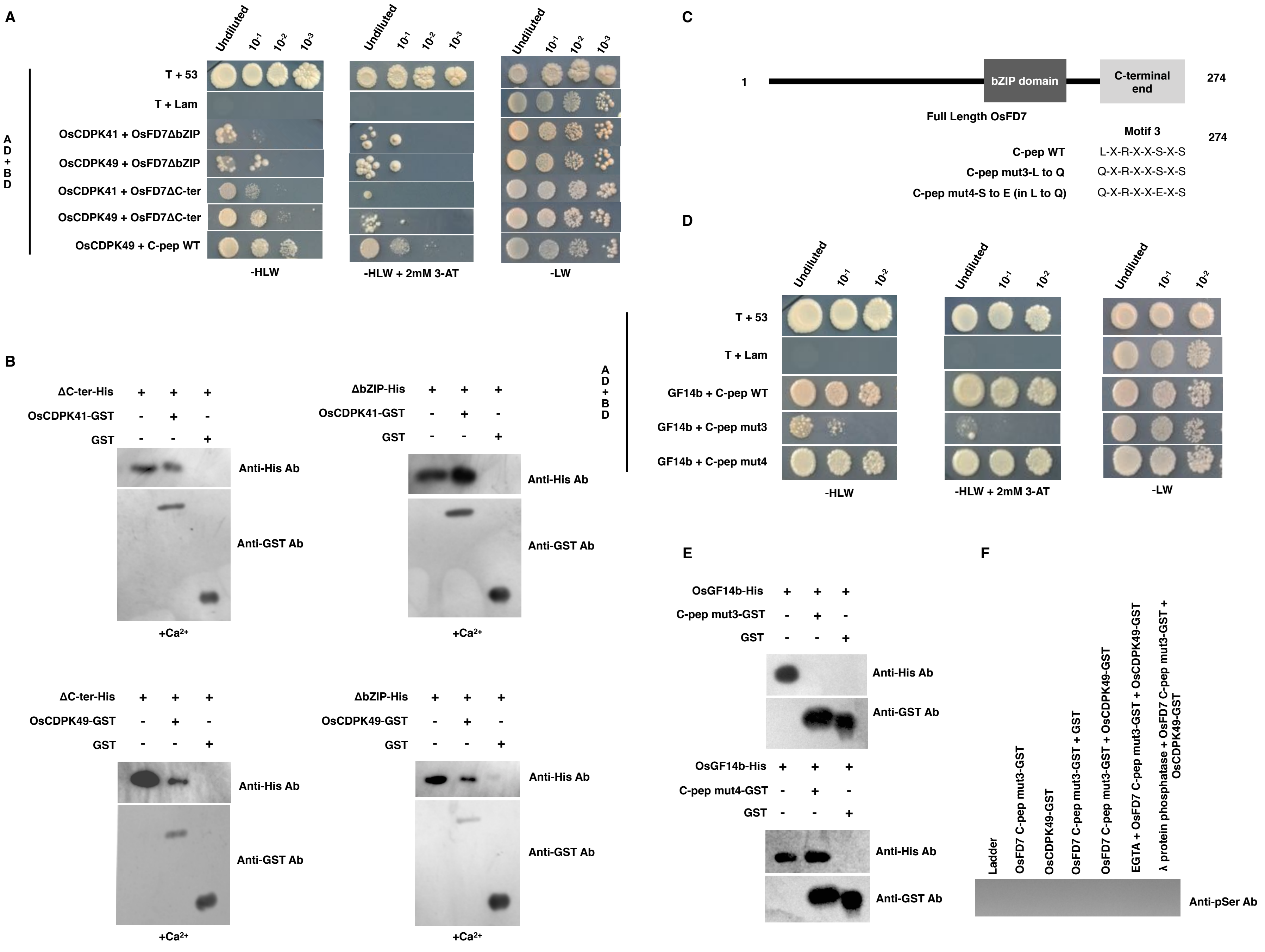


**Figure S10. Interaction of OsCDPKs and OsGF14s with truncated protein and mutated peptides of OsFD7. A:** Y2H assay depicting protein-protein interactions using truncated OsFD7. Positive control: pGADT7-T + pGBKT7-53; Negative control: pGADT7-T + pGBKT7-Lam. AD and BD denotes activation domain and DNA binding domain of GAL4, respectively. -HLW+3-AT: selective medium (SD/-His-Leu-Trp) supplemented with 3-AT; -LW: control medium (SD/-Leu-Trp). **B:** GST pull-down assay of OsCDPKs with truncated OsFD7. ΔbZIP-His: truncated OsFD7 His tagged protein; ΔC-ter-His: truncated OsFD7 His tagged protein; OsCDPK41,49-GST: GST tagged OsCDPK41,49 protein; GST: GST protein. **C:** Diagrammatic representation of mutated peptide of OsFD7 harbouring amino acid substitutions. **D:** Y2H assay using truncated OsFD7 and mutated OsFD7 peptides. Positive control: pGADT7-T + pGBKT7-53; Negative control: pGADT7-T + pGBKT7-Lam. AD and BD denotes activation domain and DNA binding domain of GAL4, respectively. -HLW+3-AT: selective medium (SD/-His-Leu-Trp) supplemented with 3-AT; -LW: control medium (SD/-Leu-Trp). **E:** GST pull-down assay performed with mutated peptides of OsFD7. C-pep mut3,mut4-GST: GST tagged mutated OsFD7 C terminal peptides; GST: GST protein. **F:** *In vitro* kinase assay using GST tagged C-terminal peptide of OsFD7 by OsCDPK49 using anti-phosphoserine. No band could be seen with mutated C-terminal peptide of OsFD7. OsFD7 C-pep mut3-GST: OsFD7 C-terminal GST tagged peptide with a single amino acid substitution LtoQ; OsCDPK49-GST: GST tagged OsCDPK49 protein; GST: GST protein.

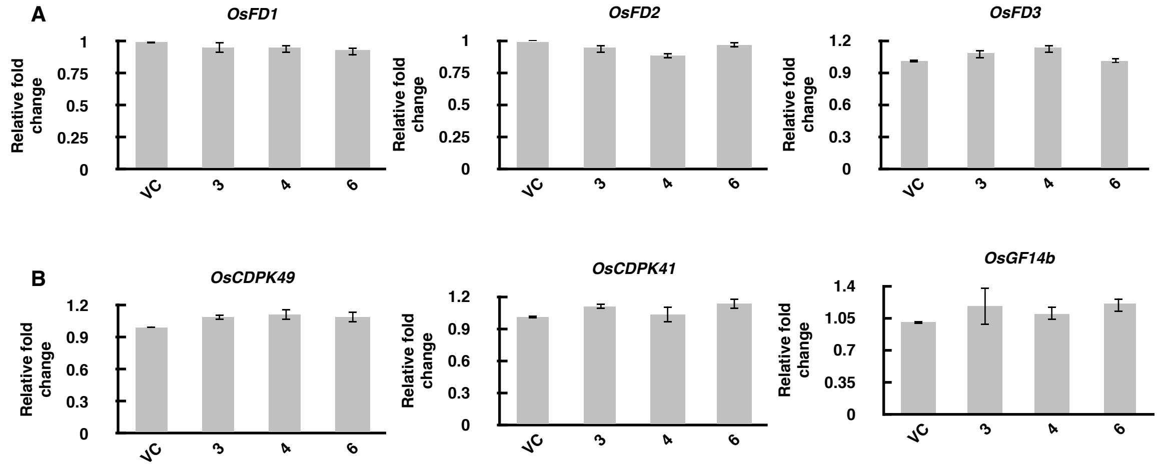


**Figure S11. Real-time PCR analysis of A:** *OsFDs and* **B:** *OsCDPK41/49 and OsGF14b* genes in *OsFD7* RNAi transgenic lines with respect to VC at SAM. *UBQ5* and *eEF-1α* genes were used as internal controls (Jain et al., 2006). Error bars represent standard error.

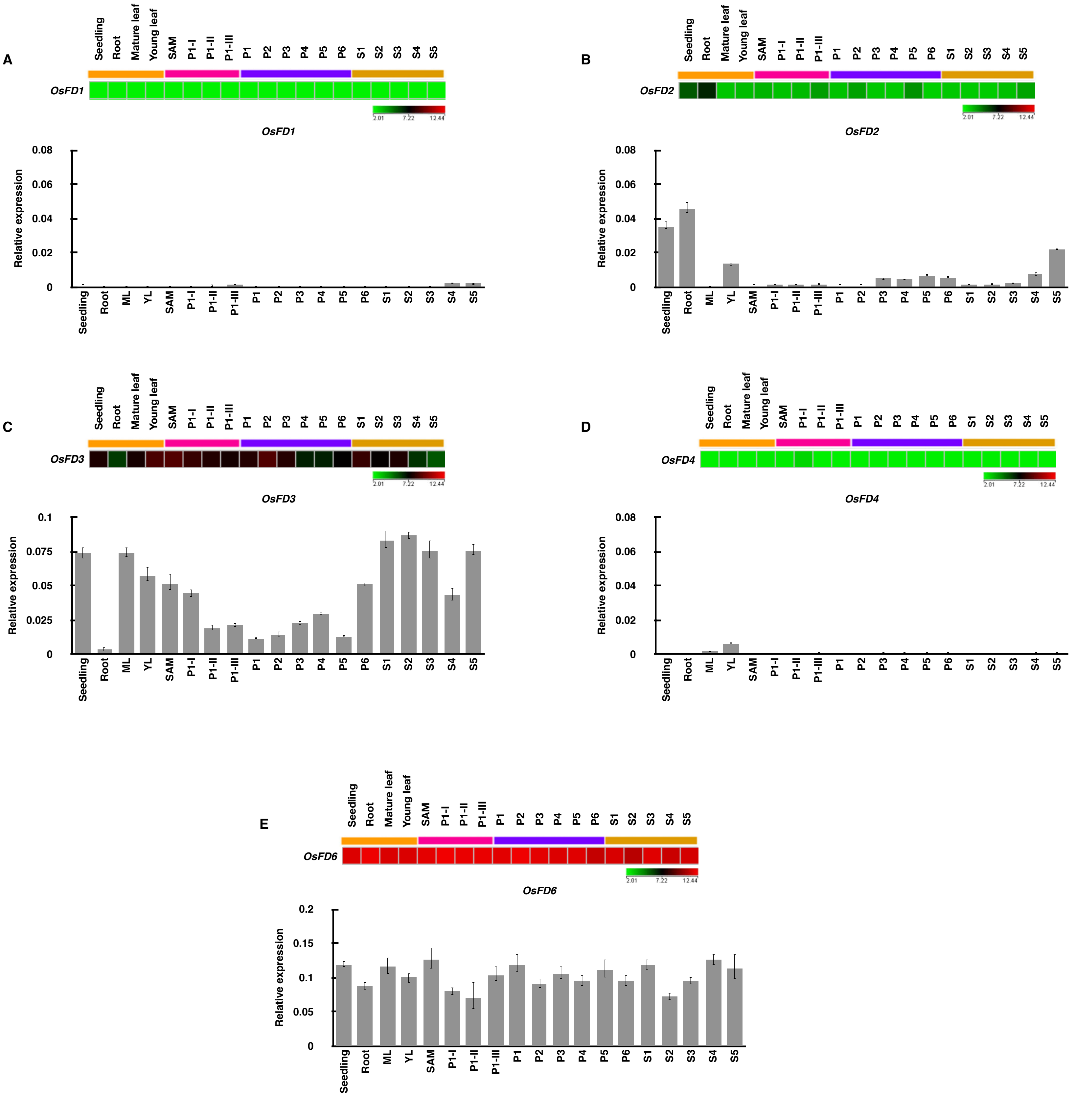


**Figure S12. Real-time PCR analysis of A:** *OsFD1*, **B:** *OsFD2*, **C:** *OsFD3*, **D:** *OsFD4*, **E:** *OsFD6 genes* in different developmental stages of rice. *UBQ5* and *eEF-1α* genes were used as internal controls (Jain et al., 2006). Error bars represent standard error.
